# Ebola virus VP35 NNLNS motif modulates viral RNA synthesis and MIB2-mediated signaling

**DOI:** 10.1101/2025.07.27.667045

**Authors:** Grace Uwase, Priya Luthra, Olivia A. Vogel, Jyoti Batra, Bruno A. La Rosa, Kathleen C. F. Sheehan, Oam Khatavkar, Jacqueline E. Payton, Robert A. Davey, Nevan J. Krogan, Christopher F. Basler, Daisy W. Leung, Gaya K. Amarasinghe

## Abstract

Ebola virus (EBOV) is a non-segmented, negative-sense virus (NNSV) with a single-stranded RNA genome. EBOV encodes for a limited number of proteins and thus depends on host factors to facilitate viral replication and pathogenesis. Of the virus-encoded proteins, multifunctional EBOV VP35 (eVP35) is necessary for host immune evasion and viral RNA synthesis. Previous proteomics studies identified an interaction between eVP35 and the host E3 ubiquitin ligase Mindbomb 2 (MIB2). Here, we show how a previously uncharacterized NNLNS motif (residues 201-205) within eVP35 serves as a binding site for MIB2. This motif is critical for eVP35-dependent inhibition of MIB2-mediated IFN induction. It is also important for EBOV RNA synthesis as MIB2 binding to eVP35 inhibited EBOV minigenome activity. Altogether, these findings highlight the importance of the eVP35 protein and the role of host factors in EBOV infection.

**SIGNIFICANCE STATEMENT:** The Ebola virus (EBOV) genome encodes for a limited number of proteins and depends on host factors to facilitate viral replication. Identification and characterization of host-viral interactions are needed to define infection, resolution, and to develop new therapeutics. EBOV VP35 (eVP35) is necessary for mediating host immune evasion and a cofactor for viral RNA synthesis. Here we characterized an interaction between eVP35 and MIB2. We show that the ^201^NNLNS^205^ motif in eVP35 is necessary and sufficient for MIB2 binding and inhibition of MIB2-mediated IFN production. Our results also reveal how the eVP35-MIB2 interaction impacts virus infection. These results support the importance of the multifunctional eVP35 to EBOV infection and highlight the significance of host proteins, including E3 ligases, during viral infection.

## INTRODUCTION

Ebola virus (EBOV), a member of the *Filoviridae* family, causes Ebola virus disease (EVD), which is associated with severe hemorrhagic fever and case fatality rates as high as 90%.^1^ Since 1976, periodic EBOV outbreaks have occurred in West and Central Africa, with the most recent outbreak ending in January 2023 in Uganda.^2–4^ These repeated outbreaks highlight the significant public health threat posed by EVD and their socioeconomic impact. Despite progress in our understanding of EBOV pathogenesis, questions remain regarding viral persistence and recrudescence ^5–7^, which underscore the ongoing need to define new therapeutic targets by delineating molecular interactions that facilitate EBOV infections.

EBOV is an enveloped filamentous virus that contains a non-segmented negative-sense single-stranded 19 kb RNA genome encoding 7 genes: nucleoprotein (NP), viral protein 35 (VP35), viral protein 40 (VP40), glycoprotein/soluble glycoprotein (GP/sGP), viral protein 30 (VP30), viral protein 24 (VP24), and large (L) polymerase.^3,8^ Of these, the EBOV VP35 (eVP35) protein, produced from the VP35 gene, is a critical cofactor for both viral genomic replication and transcription as well as for mediating viral innate immune antagonism. The eVP35 N-terminal region contains an NP binding peptide (NPBP), which regulates NP oligomerization and ssRNA binding, followed by an oligomerization domain (OD) that bridges the interaction between NP and L polymerase important for RNA synthesis (**Fig. 1A**).^9–12^ The C-terminal interferon (IFN) inhibitory domain (IID) is largely responsible for innate immune antagonism.^13^ It contains two charged patches, one that binds NP and impacts viral replication and the other that binds dsRNA to prevent pathogen-associated molecular pattern (PAMP) sensing by RIG-I and MDA5.^14–17^ Mutations that disrupt interactions with dsRNA render EBOV avirulent in mice, guinea pigs, and non-human primates, highlighting the importance of eVP35 in EBOV pathogenesis.^18–20^ eVP35 can also interact with and impair other components within the type I IFN induction and response pathways, including TBK1 and IKKε kinases as well as host adaptor protein PACT. ^8–20^ The eVP35 IID is also important for these interactions although the specific mechanisms of binding are not fully defined.^21–23^ A 65-residue conformationally flexible linker connects the eVP35 OD to IID. When bound to L polymerase, some linker residues can adopt an α-helical fold.^12,24^

**Figure 1.**
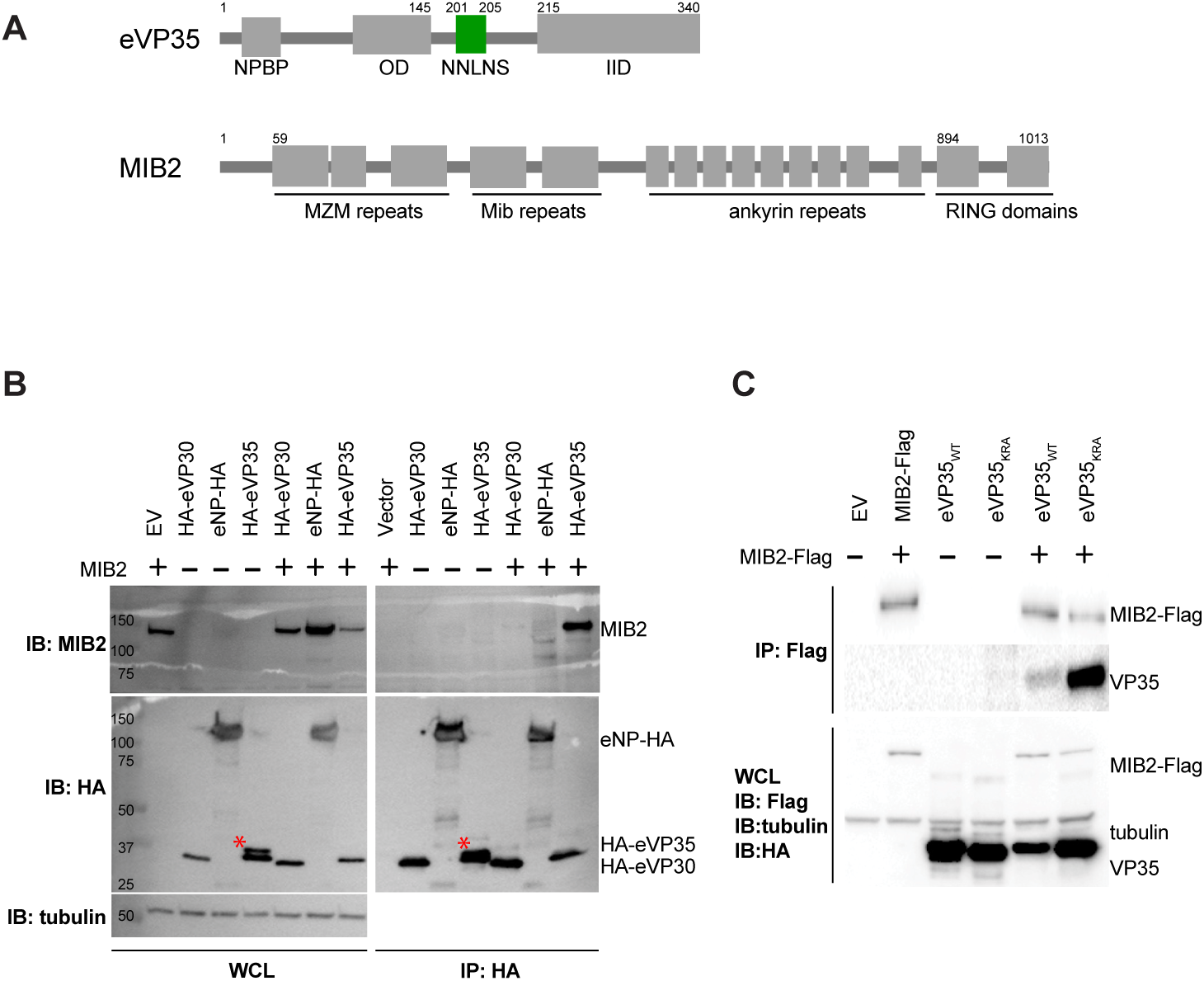
eVP35 binds MIB2. (**A**) Domain organization of full-length eVP35 and MIB2. NPBP = NP-binding peptide, OD = oligomerization domain, IID = interferon-inhibitory domain. (**B**) MIB2 interaction is specific to eVP35 but not eNP or eVP30. HA-tagged EBOV proteins were co-expressed with MIB2 for 48 h, and a co-IP experiment was performed with HA magnetic beads. Protein complexes were detected by western blotting. β-tubulin was used as a loading control. * denotes a non-specific band. (**C**) MIB2-eVP35 interaction is independent of eVP35 RNA binding ability. eVP35 WT or the eVP35 KRA mutant was co-expressed with MIB2-Flag, and co-IP with Flag magnetic beads was performed as described above. Each blot is representative of at least 2 independent replicates. For each experiment, the total amount of DNA transfected was kept constant by supplementing with empty pCAGGS vector to control for variations in transfection efficiency. EV = empty pCAGGS vector; WCL = whole cell lysate; IB = immunoblot, IP = immunoprecipitate.

Prior proteomics studies identified multiple host factors that interact with eVP35, including Mindbomb protein 2 (MIB2, also known as skeletrophin).^25,26^ Our 2018 affinity purification-mass spectrometry (AP-MS) study corroborated this finding, but MIB2 was not included in the final list of published hits due to the criteria that were used in rank ordering.^27^ However, our follow-up study also confirmed MIB2 as an eVP35 binder.^26^ MIB2 is a RING-type E3 ubiquitin ligase that regulates the NOTCH signaling pathway and has 4 identifiable domains: the N-terminal Mib-HERC2 domains that flank a ZZ-type zinc finger domain, a tandem repeat of sequences known as the Mib domains that are specific to Mindbomb proteins, a stretch of 9 ankyrin repeats, and a C-terminal RING domain containing two RING fingers with E3 ubiquitin ligase activity (**Fig 1A**).^28^ MIB2 has been shown to play an important regulatory role in innate immunity, targeting type I IFN and NFκΒ signaling pathways.^29–32^ MIB2 ubiquitinates TBK1 via K63 linkages, which is necessary for its activation and downstream phosphorylation of IRF3 and IRF7.^29^ MIB2 also enhances inflammation by promoting proteasomal degradation of CYLD, a deubiquitinase that negatively regulates various steps of both proinflammatory IRF and NF-kB pathways.^30^ CYLD downregulates IFN-β production by deubiquitinating and inactivating RIG-I and TBK1, thus inhibiting RIG-I-mediated antiviral responses.^33^ CYLD also deubiquitinates NEMO and RIPK1 to inhibit NF-κB activation, which controls the transcription of genes involved in cytokine production, cellular adhesion, inflammation, and apoptosis.^34^ While a role for MIB2 in various cancers has been described ^28,35,36^, a role in viral infection has not been well explored.

Here we characterized the interaction between MIB2 and eVP35 *in vitro* and validated the impact of MIB2 binding on eVP35 function in cell-based studies. We identify a region in the linker between the eVP35 OD and IID, named the ^201^NNLNS^205^ motif, that is important for MIB2 binding. This interaction impacts both eVP35 and MIB2 mediated functions. Recombinant EBOV containing mutations in the ^201^NNLNS^205^ motif displayed reduced cytopathic effect consistent with this region being functionally important for viral infections. Our results add to a growing body of data describing how eVP35 carries out multiple functions during infection and further highlight the significance of host factors in EBOV infection.

## RESULTS

### eVP35 binds MIB2

To validate the interaction between eVP35 and MIB2, we performed a co-immunoprecipitation (co-IP) experiment in which HA-tagged eVP35 was expressed in HEK293T cells alone or with MIB2, and immunoprecipitated with HA-magnetic beads. eNP-HA and HA-VP30 were also tested as controls. Western blot (WB) analysis revealed that MIB2 specifically co-precipitated with HA-eVP35 and not with NP-HA or HA-VP30 (**Fig. 1B**). These results confirm the proteomics results ^25,26^ and the specificity of the eVP35-MIB2 interaction.

eVP35 also binds dsRNA,^16,17^ and to eliminate the possibility that RNA contributes to the interaction between eVP35 and MIB2, we tested an eVP35 dsRNA-binding mutant containing K319A and R322A mutations (eVP35 KRA) in our MIB2 binding assays^13,17,18^. Reciprocal co-IPs with Flag-MIB2 and untagged eVP35 wildtype (WT) or eVP35 KRA showed that both eVP35 proteins co-immunoprecipitate with MIB2 (**Fig. 1C**). These results indicate that MIB2 binding to eVP35 is independent of dsRNA.

### eVP35 linker residues 201-205 are critical for MIB2 binding

We next determined whether the MIB2-eVP35 interaction is direct and defined a region in eVP35 that is important for binding MIB2. A series of eVP35 truncation constructs were generated (**Fig. 2A**), including full-length eVP35 (eVP35, residues 1-340), N-terminal region containing the NPBP and OD (eVP35_1-215_), and IID (eVP35_215-340_). eVP35 constructs were expressed as maltose binding protein (MBP) fusion proteins and tested for binding to full-length MIB2 (MIB2, residues 1-1013) using an *in vitro* pulldown assay. Our results revealed that MIB2 binds eVP35_1-340_ (**Fig. 2B**) and eVP35_1-215_ (**Fig. 2C**) but not eVP35_215-340_ (**Fig. 2D**) or the control with maltose binding protein (MBP) (**Fig. 2E**), which was also confirmed by size exclusion chromatography (SEC) experiments (**Fig. S1A**). Because MIB2 is unstable and prone to degradation and aggregation, we tested various constructs for binding to MBP-VP35_FL_ to identify the most stable construct that also maintains binding to eVP35. MIB2_FL_, MIB2_ΔRING_, and MIB2_59-894_ were tested and all maintained binding to eVP35 (**Fig. S1C**, **S1D, and Fig. S1E**) and not to the MBP control (**Fig. 1F**). We further narrowed the binding region with additional truncations of eVP35_1-215_ and tested binding to MIB2_59-894_ in *in vitro* pulldown assays and a summary of the constructs and outcomes are shown in **Fig. S2A**. We also found that the first 190 residues and the last 10 residues in eVP35_1-215_ are dispensable for MIB2 binding (**Fig. S2A**). Representative binding data for MBP-eVP35_150-205_ (**Fig. S2B**), MBP-eVP35_150-200_ (**Fig. S2C**), MBP-eVP35_150-195_ (**Fig. S2D**), and MBP-eVP35_150-190_ (**Fig. S2E**) are shown. Taken together, these results point to residues 190-205 being required for eVP35 to interact with MIB2 (**Fig. S2F**).

**Figure 2.**
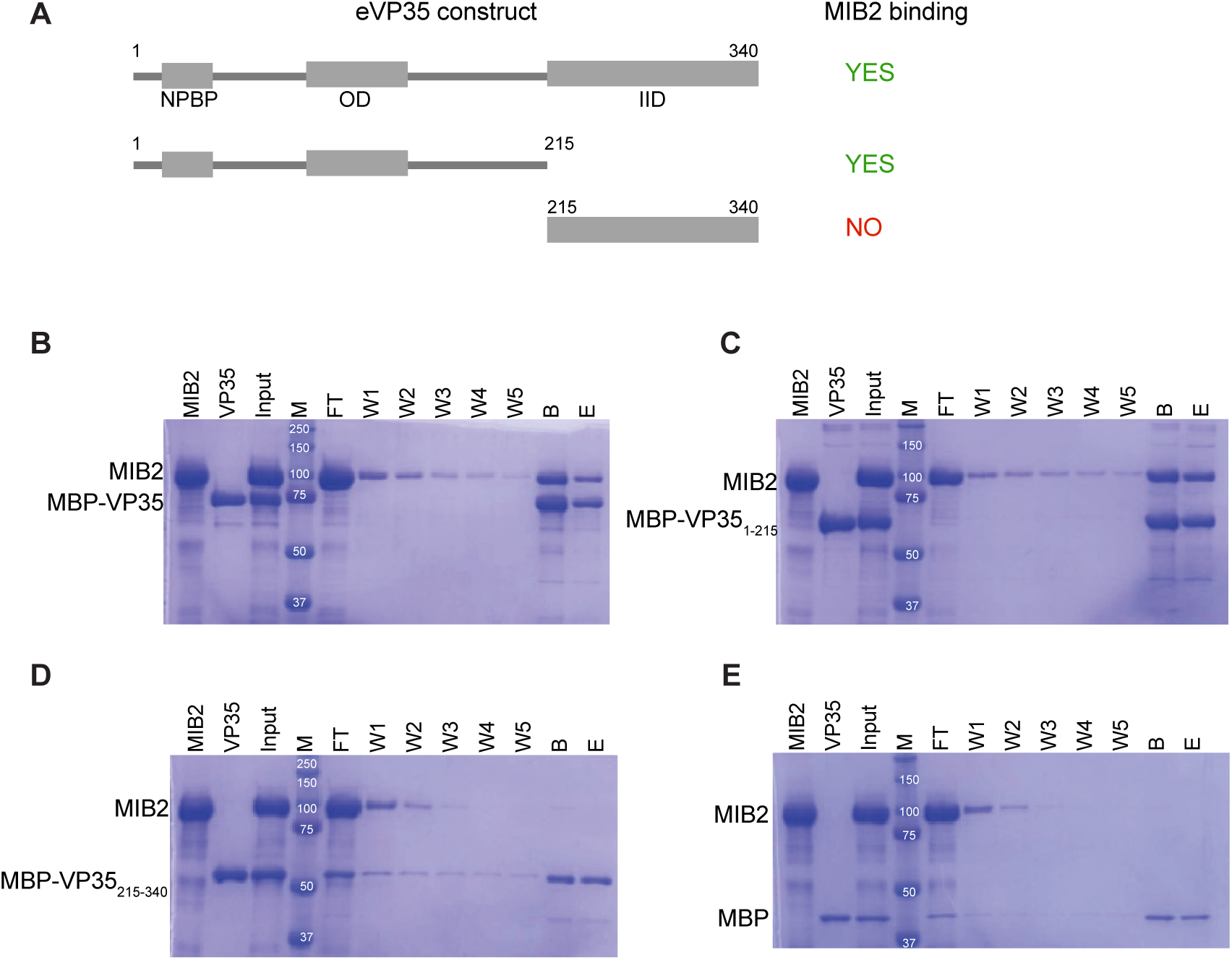
eVP35 N-terminal region mediates interaction with MIB2. (**A**) Domain architecture of critical eVP35 constructs and binding summary with MIB2.Purified MBP-eVP35 (**B**), MBP-eVP35_1-215_ (**C**), MBP-eVP35_215-340_ (**D**), or MBP (**E**) were immobilized on amylose resin prior to the addition of purified MIB2_FL_. After 5 washes, the complex was eluted with 1% maltose. Shown are the Coomassie-stained SDS-PAGE gels of each sample. Each gel is representative of at least 2 independent replicates. (Input (I), marker (M), flow-through (FT), washes (W1, W2, W3, W4, and W5), beads after washes (B), and elution (E) are indicated above each lane.

Based upon our pulldown assay results, we next used a fluorescence polarization (FP) assay to validate eVP35 binding to MIB2_59-894._ We used an eVP35 peptide encompassing residues 190-215 with a fluorescein isothiocyanate (FITC) conjugated to the N-terminus (FITC-VP35_190-215_). We found that FITC-VP35_190-215_ bound to MIB2_59-894_ with an apparent K_D_ (K_D,app_) of 5.4 ± 0.9 μM (**Fig. 3A**). This value is similar to that determined by ITC for eVP35_1-215_, indicating that the FITC label does not interfere with eVP35-MIB2 binding. We next tested the ability of shorter peptides within eVP35_190-215_ to compete and bind to MIB2_59-894_. By comparing the measured K_D,app_ values of different peptides tested, we determined that eVP35 residues ^201^NNLNS^205^ are necessary for the interaction with MIB2 (**Fig. 3B**, **Table 1**). To confirm this observation, these five residues were mutated to alanine (eVP35 5A). We note that mutation of these residues did not cause observable changes in the overall hydrodynamic properties of the protein (**Fig. S2G**). To further characterize the eVP35-MIB2 interaction, we used isothermal calorimetry (ITC) and measured a K_D_ of 5.4 ± 2.5 μM for eVP35_1-215_ binding to MIB2_59-894_ (**Fig. 3C & S2H, left panel**). ITC measurements revealed that eVP35 5A did not bind to MIB2_59-894_ (**Fig 3C, right panel**), which was further validated by an *in vitro* pulldown assay (**Fig. 3D**). Furthermore, mutation of two or three residues within this motif (eVP35_1-215_ mutants ^201^NN^202^/2A and ^203^LNS^205^/3A) resulted in near complete loss of binding to MIB2_59-894_ (**Fig. S2H**), indicating that all five eVP35 residues are critical for mediating the interaction with MIB2.

**Figure 3.**
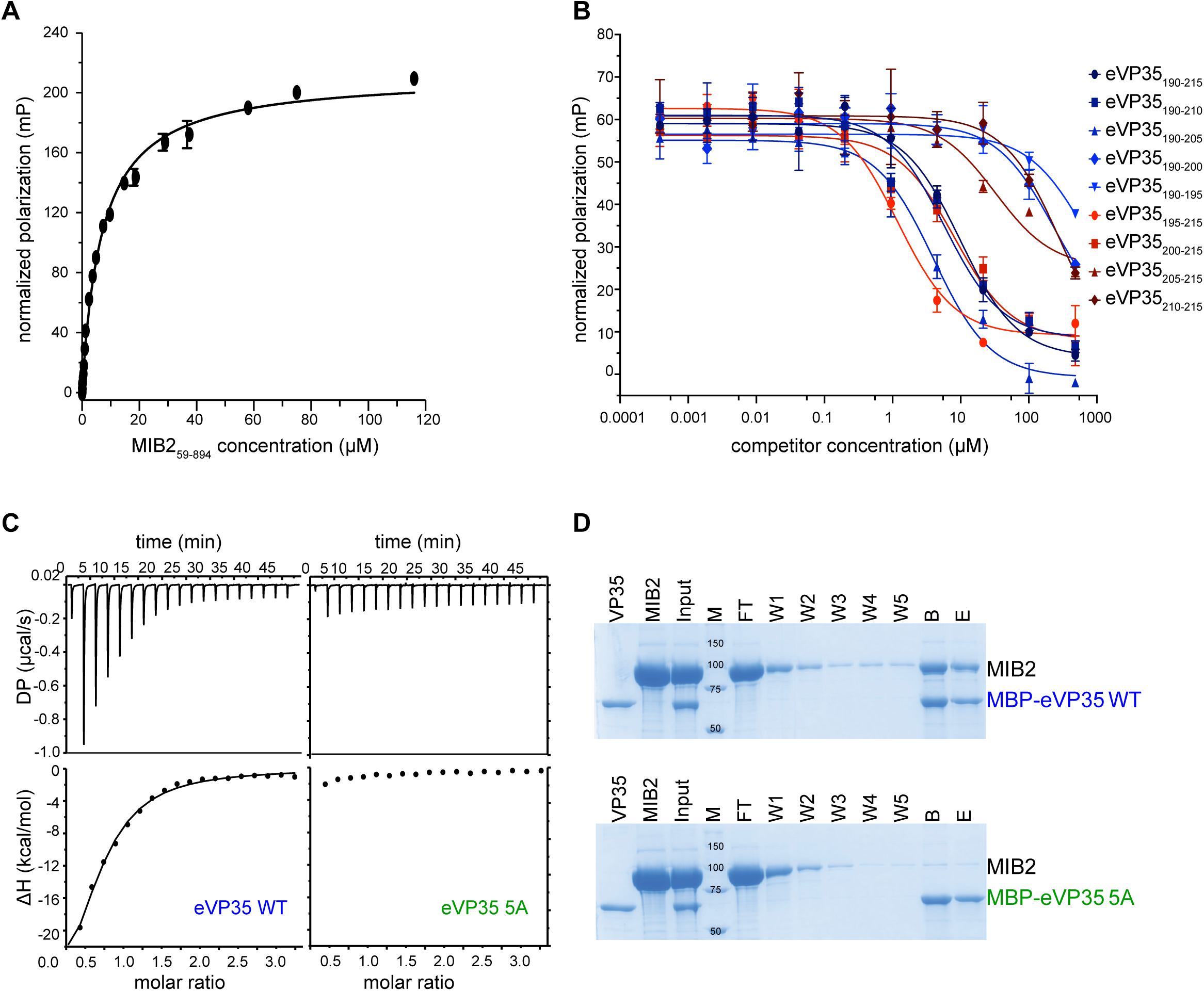
eVP35 linker residues 201-205 are critical for MIB2 binding. (**A**) Titration with MIB2_59-894_ results in an increase in fluorescence polarization of FITC-eVP35_190-215_. A K_D,app_ of 5.4 ± 0.9 μM was calculated for FITC-eVP35_190-215_ from non-linear fitting (solid line) using Prism. Error bars represent the s.d. of four independent experiments, technical duplicates each. (**B**) Competition with unlabeled eVP35 peptides with 2.5 nM FITC-eVP35_190-215_ in the presence of 2 μM MIB2_59-894_ resulted in the indicated competition curves. K_D,app_ of each peptide is summarized in Table 1. K_D,app_ of each peptide is summarized in Table 1. K_D,app_ of peptides lacking residues in ^201^NNLNS^205^ could not be detected, indicating loss of binding. Error bars represent s.d. of two independent experiments, technical duplicates each. (**C**) Representative ITC raw data and corresponding binding isotherms show a loss of eVP35 binding to MIB2 when residues ^201^NNLNS^205^ are mutated. Left panel, MIB2_59-894_ in the cell, MBP-VP35_1-215_, WT in the injection. K_D_ of 5.4 ± 2.5 μM, N = 0.8 ± 0.2. Right panel, MIB2_59-894_ in the cell, MBP-eVP35_1-215_ 5A in the injection. K_D_ = N.D, N = N.D. The data shown are representative of at least 3 independent experiments. N.D, not detected. (**D**) Pulldown showing loss of binding between MIB2 and eVP35 5A mutant. Purified MBP-eVP35_1-215_ WT (top) or MBP-eVP35_1-215_ 5A (bottom) were immobilized on amylose resin prior to the addition of purified MIB2_59-894_. After 5 washes, the complex was eluted with 1% maltose. Shown are the Coomassie-stained SDS-PAGE gels of each sample. Each gel is representative of at least 2 independent replicates. (Input (I), marker (M), flow-through (FT), washes (W1, W2, W3, W4, and W5), beads after washes (B), and elution (E) are indicated above each lane.

**Table 1:** Summary of eVP35 peptides apparent K_D_ in competition fluorescence polarization assay. N.D., not determined. app., apparent.

| SEQUENCE | $K_{D, \text{app}}$ |
| --- | --- |
| <sup>190</sup>ETVPQSVREAFNNLNSTTSLTEENFG<sup>215</sup> | $4.71 \pm 2.20$ |
| <sup>190</sup>ETVPQSVREAFNNLNSTTSLT<sup>210</sup> | $2.84 \pm 1.10$ |
| <sup>190</sup>ETVPQSVREAFNNLNS<sup>205</sup> | $1.79 \pm 0.92$ |
| <sup>190</sup>ETVPQSVREAF<sup>200</sup> | N.D. |
| <sup>190</sup>ETVPQS<sup>195</sup> | N.D. |
| <sup>195</sup>SVREAFNNLNSTTSLTEENFG<sup>215</sup> | $0.68 \pm 0.04$ |
| <sup>200</sup>FNNLNSTTSLTEENFG<sup>215</sup> | $5.14 \pm 2.73$ |
| <sup>205</sup>STTSLTEENFG<sup>215</sup> | $37 \pm 23$ |
| <sup>210</sup>TEENFG<sup>215</sup> | N.D. |

### eVP35 inhibits MIB2-mediated IFN induction

Because MIB2 promotes IFN induction and eVP35 is a potent antagonist of innate immune responses, we next investigated the functional role of the ^201^NNLNS^205^ motif in eVP35-mediated IFN inhibition. Overexpression of MIB2 alone or in combination with the RIG-I caspase activation and recruitment domain (CARD) stimulated IFN-β promoter activity (**Fig. S3A-B,** bar #3). As expected, co-expression with eVP35 WT inhibited IFN-β activity in a dose-dependent manner (**Fig. S3B,** blue bars). In contrast, the eVP35 5A mutant, which cannot bind MIB2, showed diminished inhibition of IFN-β activity (**Fig. S3B,** green bars).

MIB2 stimulates IFN production by ubiquitinating key members of the type I IFN pathway, including TBK1 kinase and CYLD deubiquitinase. We mutated eight cysteine residues (C890, C893, C921, C924, C969, C972, C998, C1001) in the C-terminal RING domains of MIB2 (MIB2 RINGm) to disrupt catalytic activity. Expression of MIB2 RINGm did not stimulate IFN-β production to the same level as MIB2 WT (**Fig. S3A & S3C**), and in the presence of MIB2 RINGm, there was no significant difference in IFN antagonism between eVP35 WT and eVP35 5A (**Fig. S3C**).

Although the eVP35 5A mutant exhibited a significant reduction in its ability to inhibit IFN induction compared to WT, this loss was only partial, which indicates the existence of another mechanism of IFN-β inhibition. eVP35 is known to inhibit IFN-β signaling by binding to double-stranded RNA (dsRNA) through the IID, thereby blocking PAMP recognition^16–18^. Previous studies have demonstrated that eVP35 mutants, which are unable to bind dsRNA, still retain some ability to inhibit virally induced type I IFN production^16,18^. This led us to hypothesize that mutations affecting both MIB2 and dsRNA binding would have an additive effect on eVP35-mediated IFN antagonism. To test this, we generated an eVP35 5A/KRA mutant that combines mutation of the ^201^NNLNS^205^ motif with mutation in the IID. The eVP35 KRA mutant alone exhibited reduced IFN antagonism compared to both eVP35 WT and the eVP35 5A mutant (**Fig. 4A, orange**). Remarkably, the eVP35 5A/KRA mutant completely lost its ability to inhibit IFN-β activity, even at the highest concentrations tested (**Fig. 4A**, cyan). Interestingly, when IFN-β was induced using CARD alone (without MIB2), the eVP35 5A mutant inhibited IFN-β activity to a similar extent as eVP35 WT (**Fig. S3D**) whereas the eVP35 KRA and 5A/KRA mutants failed to antagonize IFN-β activity (**Fig. S3D**). This suggests that while MIB2 contributes to IFN induction, its effect is less potent compared to dsRNA sensing and as a result, the impact of MIB2 inhibition is masked in the absence of MIB2 co-stimulation.

**Figure 4.**
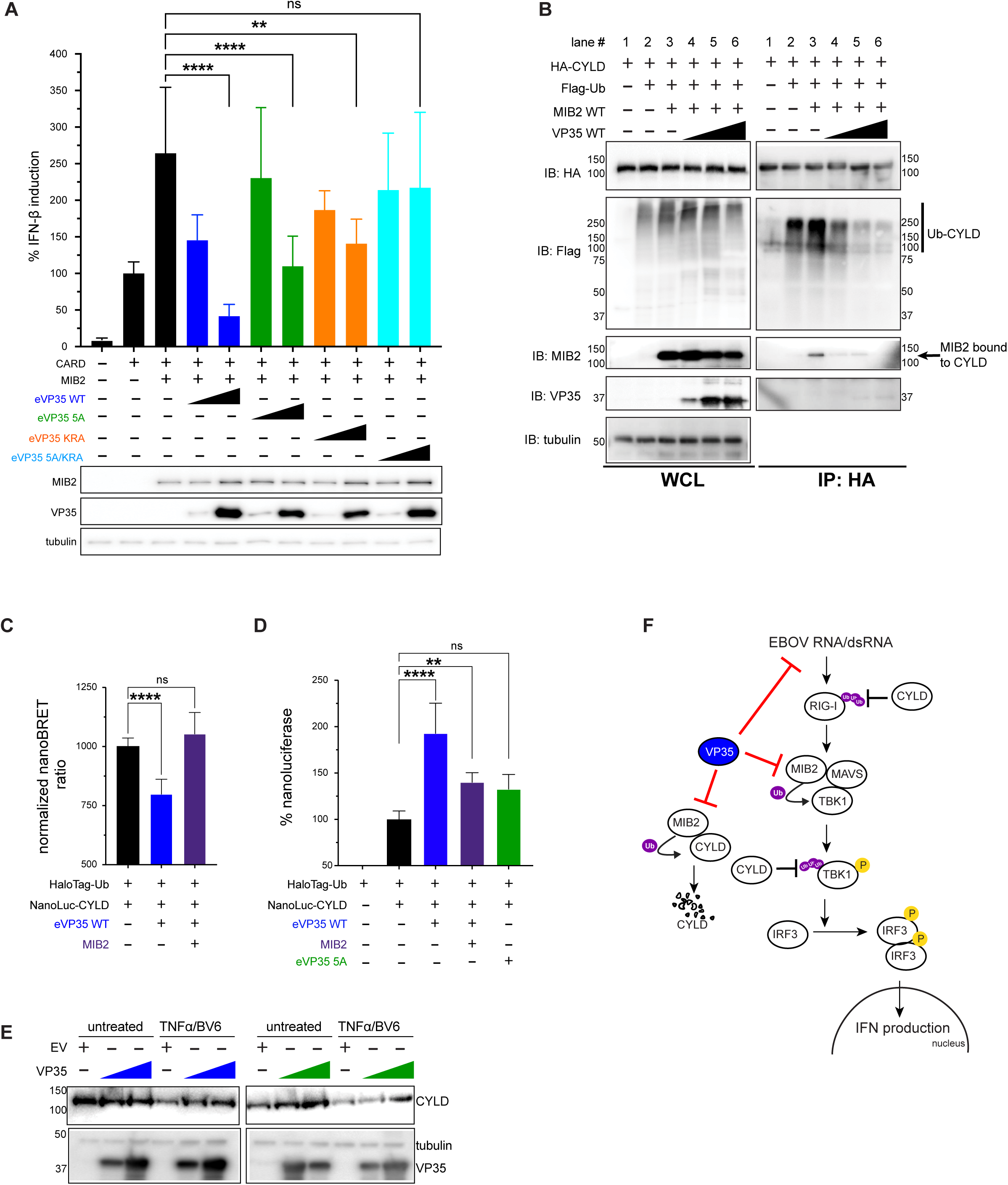
eVP35 inhibits MIB2-mediated IFN induction. (A) IFN-β promoter luciferase reporter assay when IFN production was co-stimulated with 50 ng CARD domain and 500 ng MIB2 WT (black), in the presence of 100 or 1000 ng eVP35 WT (blue), 5A (green), KRA (orange), or 5A/KRA mutant (cyan). All plasmids were transfected in HEK 293T cells, and 48 h post-transfection IFN-β activity was assessed by measuring luciferase activity. Statistical comparisons are between CARD + MIB2 and each eVP35 at the highest concentration. IFN-β induction in the presence of RIG-I CARD alone was set to 100%, and all other conditions normalized against it. Below the graph is WB validation of protein expression. The statistical significance is as indicated: not significant: n.s., *p < 0.05, **p < 0.01, ***p < 0.001, and ****p < 0.0001. (B) eVP35 inhibits MIB2-mediated CYLD ubiquitination. HA-CYLD, Flag-ubiquitin, and MIB2 were co-transfected in the presence or absence of eVP35 WT as indicated, and immunoprecipitation with HA magnetic beads was performed. WB was used to analyze ubiquitination levels of CYLD and CYLD interaction with MIB2. Ubiquitinated CYLD is indicated (**C**) NanoBRET ubiquitination assay demonstrating that eVP35 reduces CYLD ubiquitination levels. NanoLuc-CYLD (0.02 μg) and HaloTag-Ub (2 μg) were expressed in 293T cells, with or without 0.1 μg eVP35 WT or MIB2. After 24 h, transfected cells were replated into 96 well plates and treated with either HaloTag NanoBRET 618 Ligand or a DMSO control for an additional 24 h. Cells were then treated with the NanoBRET NanoGlo substrate and NanoBRET signals were measured. Each donor and acceptor readings were normalized to the HaloTag-Ub + NanoLuc-CYLD sample (set to 100%), and NanoBRET ratios were calculated as (acceptor channel/donor channel) × 1000. (**D**) Nanoluciferase assay showing that eVP35 WT increases CYLD protein expression. similar to (C), NanoLuc-CYLD and HaloTag-Ub were expressed in the presence or absence of 0.1 μg eVP35 WT, MIB2, or eVP35 5A and treated as explained in (C). Readings from the donor channel reflect CYLD protein levels. The HaloTag-Ub + NanoLuc-CYLD sample was set to 100% and the rest of the conditions are presented as a percentage of this baseline. The statistical significance is as indicated: not significant: n.s., *p < 0.05, **p < 0.01, ***p < 0.001, and ****p < 0.0001. (**E**) 293T cells were transfected with 0.5 and 2 μg of eVP35 WT (blue) or eVP35 5A (green), and 24h later were treated with 20 ng/mL TNFα and 0.5 μM BV6 for 2h. Lysates were analyzed by WB to detect CYLD protein levels. (**F**) Proposed model of how eVP35 antagonizes type I IFN production through two independent but synergistic mechanisms. The first by binding dsRNA via its IID domain and blocking PAMP recognition, and the second by binding MIB2 via the NNLNS motif and inhibiting its ability to upregulate IFN production.

MIB2 ubiquitinates CYLD and targets it for degradation ^30^, and we tested whether eVP35 affects CYLD ubiquitination and protein levels. We co-expressed Flag-ubiquitin, HA-CYLD, and MIB2 with or without eVP35 in 293T cells. In the presence of eVP35, the CYLD-MIB2 interaction was reduced. We also noted a reduction in CYLD ubiquitination in an eVP35 dose-dependent manner (**Fig. 4B** IP lanes 4, 5, and 6). To validate and quantify the impact of eVP35 on CYLD ubiquitination, we used a NanoBRET assay by co-expressing NanoLuc-CYLD with Halo-ubiquitin. When CYLD is ubiquitinated, the proximity of NanoLuc and HaloTag generates a NanoBRET signal proportional to the extent of ubiquitination. We found that co-expression with eVP35 significantly reduced CYLD ubiquitination, whereas co-expression of eVP35 and MIB2 restored ubiquitination (**Fig. 4C**).

We assessed CYLD protein levels by expressing NanoLuc-CYLD and measuring the luminescent signal in the presence of a NanoGlo substrate. Expression of eVP35 WT significantly increased CYLD levels, suggesting that it inhibits CYLD degradation. When MIB2 was co-expressed with eVP35 WT, CYLD levels decreased (**Fig. 4D**). In contrast, the eVP35 5A mutant did not affect CYLD levels, indicating that the impact of eVP35 on CYLD degradation is mediated through MIB2 binding. A previous report showed that CYLD levels decrease when TNFα-stimulated cells are infected with Sendai virus,^33^ and we observed that treating 293T cells with a combination of TNFα and BV6 (a potent and specific antagonist of inhibitors of apoptosis proteins) induced CYLD degradation (**Fig. S3E**). eVP35 WT inhibited this degradation, whereas eVP35 5A did not (**Fig. 4E**)

Collectively, these findings demonstrate that eVP35 antagonizes IFN production through two independent mechanisms (**Fig. 4F**). First, eVP35 binds dsRNA via its IID domain and blocks PAMP recognition, as previously established ^16–18^. Second, eVP35 binds MIB2 via the ^201^NNLNS^205^ motif and inhibits MIB2-mediated IFN production by ubiquitinating CLYD for degradation. These two mechanisms are likely additive.

### eVP35 ^201^NNLNS^205^ motif contributes to EBOV RNA synthesis

Because eVP35 is a cofactor for the EBOV replication complex, we next tested the role of the ^201^NNLNS^205^ motif in viral RNA synthesis using a minigenome (MG) assay ^37,38^. Our results showed that the eVP35 5A mutant had reduced MG activity compared to eVP35 WT, indicating that these residues contribute to viral RNA synthesis (**Fig. 5A**). We also leveraged the fact that eVP35 oligomerizes into a tetramer for RNA synthesis^12,24^ and transfected cells with different ratios of eVP35 WT and eVP35 5A plasmids that would allow both versions to incorporate into a tetramer. We found that as the proportion of eVP35 5A increased relative to eVP35 WT, MG activity showed a corresponding decrease (**Fig. S4A**), indicating that the ^201^NNLNS^205^ motif also plays a critical role in viral RNA synthesis. To further test the role of eVP35 ^201^NNLNS^205^ motif in EBOV infection, we recovered a virus containing the eVP35 5A mutation and tested its ability to replicate in 293T cells. At 20 hpi, the number of infected cells was significantly higher in EBOV WT -infected cells compared to those infected EBOV containing the eVP35 5A mutant (**Fig. S4B**). This result is consistent with our minigenome data and suggests that the eVP35 5A mutation reduces replication efficiency in 293T cells. We then tested the impact of the eVP35 5A mutation on cytotoxicity using Vero E6 cells. Seven days post-infection, EBOV carrying the eVP35 5A mutation was found to cause less cell death compared to the EBOV WT virus. Reduced cytopathic effects support our minigenome experiments and suggest that these residues are indeed important for viral replication (**Figure 5B**). We also conducted quantitative viral growth curve analysis in Vero E6 cells. The resulting data, show no significant differences in viral titers between EBOV WT and eVP35 5A mutant viruses (**Fig. S4C**).

**Figure 5.**
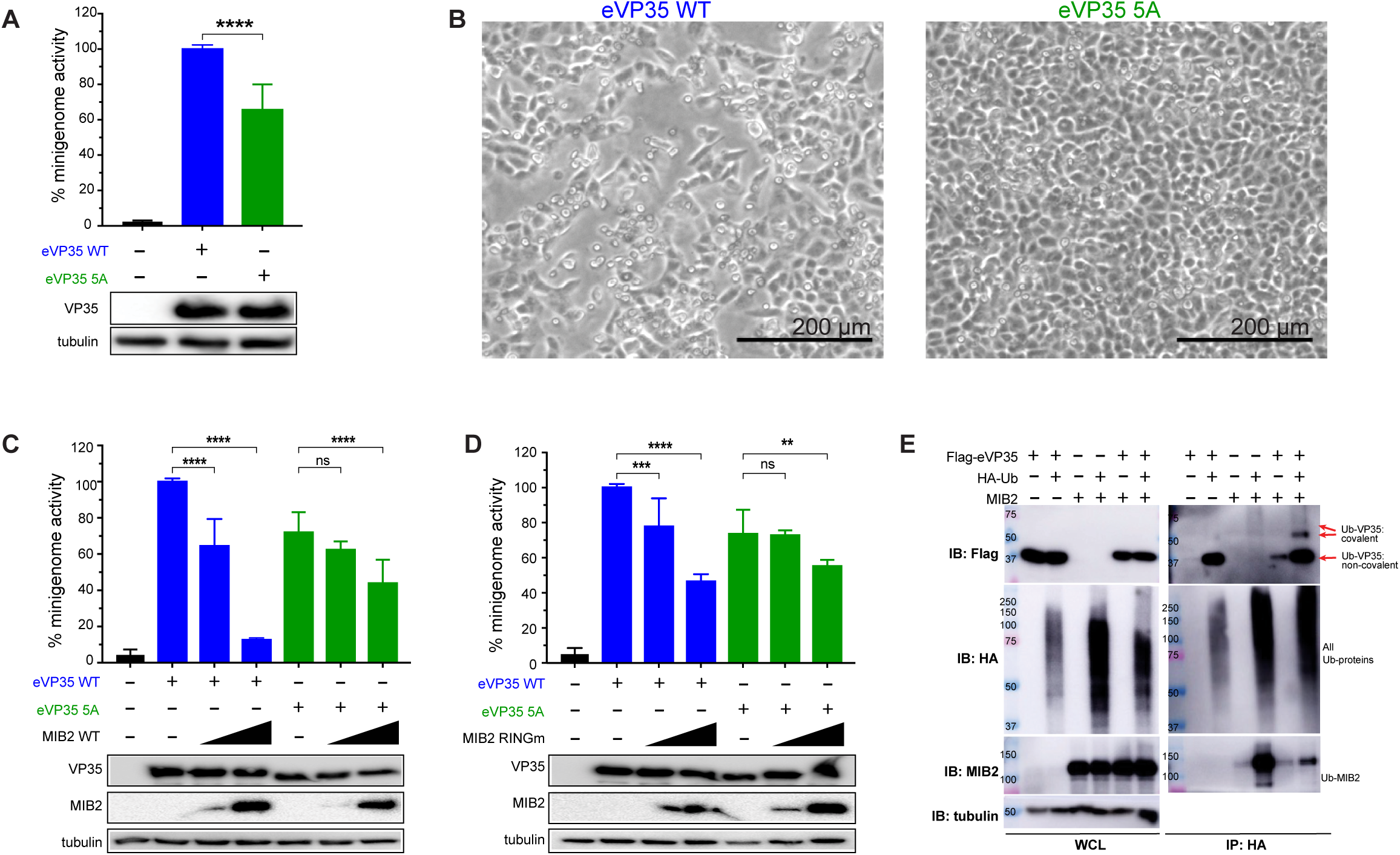
eVP35 linker residues 201-205 contribute to EBOV RNA synthesis. **(A)** eVP35 WT (blue) or 5A mutant (green) were cotransfected with plasmids required to reconstitute the EBOV RNA polymerase complex (L, NP, VP30) along with a plasmid encoding the Renilla luciferase minigenome RNA and a firefly luciferase expression plasmid, which served as a control for transfection efficiency. Relative activity was determined by setting eVP35 WT Renilla luciferase to 100% activity and normalizing the rest of the conditions against it. (**B**) Cytopathic effect of recombinantly produced EBOV with WT eVP35 or recombinant virus encoding the eVP35 5A mutation, 7 days post-infection in Vero E6 cells. (**C**) Increasing amounts of MIB2 WT plasmid (125, 500 ng) were co-transfected with the plasmids required to reconstitute the EBOV RNA polymerase complex, either with WT eVP35 (blue) or 5A mutant (green), along with a plasmid encoding the Renilla luciferase minigenome RNA and a firefly luciferase expression plasmid, which served as a control for transfection efficiency. Relative activity was determined by setting eVP35 WT Renilla luciferase to 100% activity and normalizing the rest of the conditions against it. (**D**) An increasing amount of MIB2 RINGm was added in the minigenome assay, either with WT eVP35 WT (blue) or 5A mutant (green) similar to (**A**). Protein expression was validated by WB. The statistical significance is as indicated: not significant, n.s.; *p < 0.05, **p < 0.01, ***p < 0.001, and ****p < 0.0001. (**E**) eVP35 is ubiquitinated in the presence of MIB2 WT. Flag-eVP35 and HA-Ubiquitin were transfected in the presence or absence of MIB2, and immunoprecipitation with HA magnetic beads was performed. WB was used to analyze the interaction. Ubiquitinated eVP35 is indicated with arrows.

The linker region of eVP35 forms a long loop structure over the entrance of the L polymerase active site (**Fig. S5A**) and contributes to eVP35-L interaction. Given this, we tested whether MIB2 binding to eVP35 might affect EBOV polymerase activity. Co-expression of MIB2 WT with eVP35 WT resulted in a dose-dependent inhibition of MG activity. As a control, we tested MIB1 (a homolog of MIB2) and found that it did not inhibit EBOV MG activity (**Fig. S4D**). When we co-expressed MIB2 WT with the eVP35 5A mutant, MG activity was also inhibited, but to a lesser extent (**Fig. 5C**). This suggests that the ability of MIB2 to inhibit MG activity is partially dependent on its interaction with ^201^NNLNS^205^ residues of eVP35.

Given the role of MIB2 as an E3 ligase, we next examined the MIB2 RINGm mutant. MIB2 RINGm had reduced inhibition of MG activity in the presence of eVP35 WT compared to MIB2 WT (**Fig. 5D, green bars, & S4E**). In the context of the eVP35 5A mutant, there was no difference between MIB2 WT and MIB2 RINGm (**Fig. S4E, green bars**). The difference between MIB2 WT and MIB2 RINGm in inhibiting MG activity suggests a potential role for MIB2-mediated ubiquitination in modulating the eVP35 function.

### MIB2-mediated ubiquitination of eVP35

We next asked whether eVP35 functions as a substrate for MIB2. HA-ubiquitin was co-expressed with eVP35 WT in HEK 293T cells, both in the presence and absence of MIB2. Co-IP experiments revealed that eVP35 WT is ubiquitinated in the presence of MIB2, as evidenced by the appearance of a higher molecular weight band near the 50 kDa marker that corresponds to a ubiquitinated eVP35 (**Fig. 5E**). In contrast, neither MIB2 RINGm and MIB2 ΔRING were able to ubiquitinate eVP35 WT (**Fig. S4F**). We also tested eVP35 5A and the resulting data suggests that eVP35 5A can be ubiquitinated by MIB2 (**Fig. S4F**), suggesting that direct interaction via the ^201^NNLNS^205^ region is not required for eVP35 ubiquitination. Our data also suggest that eVP35 may be associated with non-covalently linked ubiquitin, independent of MIB2, and additional studies are needed to explore this possibility.

Specific interactions between eVP35 and EBOV L protein from the structure (PDB 8JSM) is shown in **Fig. S5A**. ^12,24^ Since VP35 is conserved across multiple filoviruses, we also tested whether VP35 from other filoviruses can interact with MIB2 by co-IP. A VP35 homolog found in the *Myotis myotis* bat genome was also tested. We found that in addition to VP35 from EBOV 1976 and EBOV 2014, VP35 encoded by the Marburg virus (MARV) and by *Myotis myotis* bat bind MIB2, suggesting that this interaction is conserved across the filovirus family (**Fig. S5B**). Further *in vitro* pull-down assays revealed that MARV VP35 does not directly bind MIB2 unlike eVP35 (**Fig. S5C & S5D**) or the purified proteins exhibit different interactions compared to those in mammalian expression. This was further confirmed by competition FP, where the eVP35 peptide (190-215) successfully competed with the FITC-labeled eVP35-MIB2 complex, while mVP35 (179-204) did not (**Fig. S5E**). Sequences corresponding to the ^201^NNLNS^205^ region of multiple VP35 proteins, including MARV, revealed that this region is not conserved in MARV (**Fig. S5F**). While the overall effect of MIB2 on filoviruses may be significant, MIB2 interaction via the ^201^NNLNS^205^motif and the corresponding functional implications may be limited to Ebola virus.

## DISCUSSION

Host-virus interactions are critical in shaping the outcome of infections. Viruses like EBOV have low coding capacity (7-10 gene products are produced during infection) and therefore encode for multifunctional proteins and utilize host factors that can facilitate immune evasion and viral replication. In this study, we characterized a novel interaction between eVP35 and the host E3 ligase MIB2 and determined its functional impact *in vitro*. eVP35 is composed of an N-terminal region and a C-terminal domain connected by a linker.^9,10,12,15,17^ Due to the intrinsic disorder and flexibility of the linker, this region has largely eluded biochemical and structural characterization. More recently, the structure of eVP35 bound to the L polymerase revealed that the linker region is conformationally labile and can become structured in the presence of a binding partner.^12^ Our studies here showed that eVP35 residues ^201^NNLNS^205^ within the linker are critical for binding to MIB2. This finding was supported by mutagenesis experiments where MIB2 binding is lost with the eVP35 5A mutant. Additionally, loss of binding was observed with ^201^NN^202^/2A and ^203^LNS^205^/3A mutants, which emphasizes the collective importance of all five residues in facilitating the eVP35-MIB2 interaction.

Prior studies established a role for MIB2 in upregulating type I IFN production by ubiquitinating key members in this pathway, including CYLD.^29,30^ Our findings support this observation, showing that overexpression of MIB2 increases IFN activity whereas an enzymatically inactive MIB2 fails to stimulate IFN activity (**Fig. S3A**). When co-expressed with the RIG-I CARD domain, MIB2 enhances IFN activity up to 6-fold compared to CARD alone (**Fig. 4A & S3B**). We show that co-expression with eVP35 WT leads to a dose-dependent loss of IFN production and that the eVP35 5A mutant has diminished ability to inhibit IFN production (**Fig. 4A & S3B**). The loss of IFN antagonism by eVP35 5A was only partial, which is consistent with the presence of other IFN antagonism mechanisms. Our studies show that an eVP35 mutant that combines the ^201^NNLNS^205^ motif and IID mutations (eVP35 5A/KRA) completely loses the ability to inhibit IFN-β activity, even at the highest concentrations tested (**Fig. 4A, cyan**). The additive effect of these mutations suggests that both mechanisms function independently of each other. Interestingly, when IFN-β was induced using CARD alone (without MIB2), eVP35 5A inhibited IFN-β activity to a similar extent as eVP35 WT (**Fig. S3D)**. In contrast, eVP35 KRA and eVP35 5A/KRA mutants failed to antagonize IFN-β activity (**Fig. S3D)**. This suggests that while MIB2 contributes to IFN induction, its effect is less potent compared to dsRNA sensing by the IID and, as a result, the impact of MIB2 inhibition is masked in the absence of MIB2 co-stimulation.

The mechanism by which eVP35 inhibits MIB2-mediated IFN production was evaluated using CYLD as a representative MIB2 substrate^30,33^. We show that eVP35 is associated with reduced CYLD ubiquitination, resulting in increased CYLD protein levels (**Fig. 4B-4E**). Notably, the eVP35 5A mutant did not affect CYLD levels, indicating that the impact of eVP35 on CYLD depends on the ability of eVP35 to bind MIB2. This finding demonstrates that eVP35 binding directly disrupts the ubiquitination process mediated by MIB2. Since MIB2 also targets substrates in other critical pathways, such as NF-κB signaling and cell death regulation, it will be interesting to determine whether eVP35’s inhibitory effect extends to these substrates or is confined to the IFN response pathway Our functional characterization also identified a role for the eVP35 ^201^NNLNS^205^ motif in filoviral RNA synthesis. Mutations of these residues exhibit a partial loss of activity in EBOV RNA synthesis (30-50% loss) as evidenced by our minigenome assays, suggesting that they also contribute to eVP35 polymerase co-factor function (**Fig. 5A & S4E**). The linker region of only one of the four eVP35 subunits was captured bound to the L polymerase (**Fig. S5**)^12^, and it was found to form a long loop structure atop the entrance of the active site of L polymerase and is stabilized by eVP35 residues L209, E211, and G215 that directly interact with L polymerase. While the ^201^NNLNS^205^ motif within this structure appears to form part of an alpha helix, the singular nature of this observation precludes definitive conclusions regarding its specificity to the interaction with L. Nevertheless, our data suggest that these residues likely adopt a stabilizing structural conformation crucial for eVP35 function in RNA synthesis. Further structural studies are necessary to elucidate the conformational dynamics of this eVP35 region and its implications for EBOV RNA synthesis. Our study also demonstrates that MIB2 inhibits EBOV polymerase activity in a dose-dependent manner, while the enzymatically inactive MIB2 has reduced inhibition (**Fig. 5C & D**). Interestingly, MIB2 WT and MIB2 RINGm have similar inhibitory activity at lower concentrations, while MIB2 RINGm has significantly reduced inhibition at higher concentrations. Since both MIB2 WT and RINGm can bind eVP35, these observations suggest that MIB2-mediated inhibition of eVP35 polymerase co-factor function involves both direct protein-protein interaction and MIB2 E3 ligase activity, underscoring the multifaceted nature of this regulatory mechanism. This is supported by the fact that in the eVP35 5A mutant, there is no difference between MIB2 WT and MIB2 RINGm in the ability to inhibit eVP35.

Given that neither version of MIB2 can directly bind eVP35 5A, the residual inhibition of minigenome activity is likely independent of the eVP35-MIB2 interaction, and further studies will be needed to elucidate this. Additionally, the observed ubiquitination of eVP35 in the presence of MIB2 supports the notion of post-translational modifications as crucial regulatory events in filovirus replication and pathogenesis. Indeed, one study has shown eVP35 ubiquitination by E3 ligase TRIM6, which has a role in promoting viral replication.^39,40^ We noticed that VP35 levels appear to decrease in the presence of MIB2 (**Fig. 1B & 1, 5E**), and it is plausible that MIB2 may ubiquitinate eVP35 and target it for degradation. Additional studies will be needed to fully evaluate the impact of ubiquitination on eVP35 function. Given that the ^201^NNLNS^205^ motif contributes to EBOV polymerase activity and is essential for interacting with MIB2, our study suggests a potential mechanism of inhibition, whereby MIB2 binding to eVP35 inhibits its ability to bind L polymerase. Furthermore, we show that recombinant EBOV containing eVP35 5A mutation results in fewer numbers of infected cells in 293T cells and reduced cytopathic effects in Vero E6 cells (**Fig. S4B, Fig. 5C**). In Vero E6 cells however, there was no difference in viral titers between EBOV WT and the 5A virus, indicating that the CPE differences are likely due to perturbations in MIB2-related signaling. MIB2 has been demonstrated to inhibit cell death by modulating the activity of RIPK1 and cFLIP in cell death pathways^32,41^, and it is possible that VP35 binds MIB2 and alters its activity in these signaling pathways similar to what we observed in IFN production. Future studies using MIB2 knockout models or reconstitution systems will be necessary to test the direct impact of VP35-MIB2 interactions on host cell death responses.

Together, our findings reveal an additional regulatory layer of eVP35. The ^201^NNLNS^205^ motif mediates critical host-pathogen interactions during infection through MIB2. Ubiquitination of eVP35, including potential MIB2-mediated ubiquitination, may further regulate eVP35 function, and ubiquitination-related studies are part of our ongoing investigation. Results here highlight the significance of the identification of new binding interfaces as they will not only deepen our understanding of viral pathogenesis but also uncover potential targets for therapeutic intervention. Given the role of post-translational modifications in acute and persistent infections, our findings may have additional implications beyond those described here. While these in vitro studies provide key insights into a role for ubiquitin ligases in EBOV infection, the significance of these findings in vivo, including those in MIB2 sufficient and deficient models, remains to be examined.

## MATERIALS AND METHODS

### Antibodies

The following antibodies were used in our study: polyclonal rabbit anti-skeletrophin/MIB2 (Abcam, ab99378), polyclonal rabbit anti-flag antibody (Sigma Aldrich, F7425), polyclonal rabbit anti-HA antibody (Sigma Aldrich, H6908), monoclonal mouse anti-β-tubulin antibody (Sigma Aldrich, T8328), monoclonal rabbit anti-CYLD (cell signaling, 8462S), and EBOV VP35 mAb (2A11.F12.D7) developed in house.

### Cell lines and viruses

HEK293T (human embryonic kidney SV40 Tag transformed; ATCC, CRL-3216) and Vero (ATCC, CCL-81) were maintained in Dulbecco’s modified Eagle’s medium (Corning Cellgro or ThermoFisher scientific) with 10% fetal calf serum (GIBCO or Gemini Bio-Products) at 37°C in a humidified atmosphere with 5% CO_2_.

A circular polymerase extension reaction (CPER) was adapted from Amarilla *et. al*.^42^ to generate recombinant Ebola virus (rEBOV, based on Genbank AF086833.2). Six EBOV fragments with 40-50 bp overlaps were amplified from a plasmid encoding the entire virus genome (p15AK EBOV Mayinga). Each fragment was purified out of an agarose gel and 0.05 pmol was mixed with a linker fragment in a 50 uL reaction with 200 µM of each dNTP, 1x PS GXL reaction buffer, and 2 μL of PrimeStar GXL DNA polymerase (Takara Bio, #R050A). The following cycling conditions were used for CPER: initial denaturation at 98 °C for 2 min, followed by 20 cycles of denaturation at 98 °C for 10 s, annealing at 55 °C for 15 s, and extension at 68 °C for 25 min, followed by a final extension at 68 °C for 25 min. For mutant incorporation, site directed mutagenesis was performed on the relevant genomic fragment by PCR. The rEBOV genome was made by substitution of the altered fragment for the wild type fragment and assembled into a full genome by CPER. To recover recombinant virus, HEK293T cells grown in a 12 well plate, were transfected with 25 uL of CPER product, and support plasmids: 0.5 ug pCAGGS NP, 0.05 ug pCAGGS VP30, 0.05 ug pCAGGS eVP35, and 0.5 ug pCAGGS L. After 4 days the supernatant was clarified, and virus was passaged twice on Vero E6 cells. A sample of culture supernatant from the second passage was collected in TRIzol LS and used to prepare RNA. RNAseq was performed on this RNA by Illumina deep sequencing. It was found that the WT and 5A mutant viruses represented >99% of recovered sequences indicating that each was stable in the virus population. Each was then used for evaluation of mutants for CPE and sensitivity to interferon.

### Plasmids

For mammalian expression: The coding sequence of human MIB2 (transcript BAB92950) cloned in pCAGGS vector was obtained from Chris Basler lab. Isoform 1 (NM_080875.5), different domains of MIB2, and the residue mutants were all cloned using transcript BAB92950 as a template. The coding sequences for EBOV proteins (VP35, VP30, NP, and L) from *Zaire ebolavirus* (NC_002549.1) were used. All MIB2 and EBOV protein inserts were cloned into NotI/XhoI sites of the pCAGGS plasmid (Addgene) by restriction digest for mammalian expression. Expression of MARV and Myotis bat VP35 plasmids in the pCAGGS backbone have been previously described. ^43,44^

For bacterial expression and *in vitro* purification, human MIB2 (transcript BAB92950) and MIB2 truncations or EBOV VP35 and eVP35 truncations were subcloned into a pET15b or a modified pET15 vector with maltose binding protein (MBP) fusion tag and pMALc vector.

### Co-immunoprecipitation Assays

HEK293T cells were transfected with the mammalian expression constructs as indicated using lipofectamine 2000 (Invitrogen) according to the manufacturer’s instructions. Cells were harvested in Tritonx-100 buffer (25 mM Tris (pH 8), 150 mM NaCl, 1 mM EDTA, 1% TritonX-100, 10% glycerol) supplemented with protease inhibitors (benzamidine, leupeptin, pepstatin A, antipain). Clarified cell lysates were incubated with anti-HA or flag-magnetic beads (Sigma-Aldrich) for 1-2 h at 4℃ followed by 3 washes with wash buffer (1x TBS (20 mM Tris, pH 7.5, 150 mM NaCl) with 0.05% triton-100). Protein complexes were eluted from the beads by direct incubation in 2X laemmli loading dye. Eluates and whole-cell lysates were analyzed by immunoblotting with the antibodies as indicated.

### Protein Expression and *in vitro* Purification

eVP35 and MIB2 (full-length proteins or truncations) were expressed as MBP fusion proteins in BL21(DE3) *E. coli* cells in Luria Broth media. Protein expression was induced at an OD600 (optical density at 600 nm) of 0.6 – 0.8 with 0.5 mM isopropylthiogalactoside (IPTG) and continued for 15 h at 18 ℃ for eVP35 or 16 ℃ for MIB2. Cells were then harvested and suspended in lysis buffer (20 mM Tris (pH 7.8), 1 M NaCl, 20 mM Imidazole, 5% glycerol, protease inhibitors, 5 mM 2-mercaptoethanol (β-ME) for VP35 constructs or including 200 mM arginine for MIB2 constructs). Suspended cells were further lysed using an EmulsiFlex-C5 homogenizer (Avestin) and clarified by centrifugation at 24,000 x g at 4 ℃ for 40 min. MBP-fusion proteins were purified using a series of affinity and ion exchange chromatography columns and tag removed prior to size-exclusion chromatography. The purity of the samples was determined by SDS-PAGE at each stage.

### In vitro pulldown assays

Amylose resin (NEB) was equilibrated with buffer containing 20 mM Tris (pH 7.5), 200 mM NaCl, and 5% glycerol. The equilibrated resin was incubated with bait protein (MBP-tagged MIB2 or eVP35) for 10 min, washed 5 times, and resuspended in buffer. Untagged prey protein (MIB2 or eVP35) was then applied to immobilized bait and allowed to incubate for 10-30 min, then complex washed for 5 times. The complex was eluted with wash buffer containing 1% maltose. Gel samples were taken at each step and visualized on SDS-PAGE by Coomassie blue staining.

### Isothermal Titration Calorimetry

eVP35/MIB2 binding assay was performed on a VP-isothermal titration calorimeter (VP-ITC) (MicroCal). Protein samples were dialyzed against buffer containing 20 mM HEPES pH 7.0, 200 mM NaCl, and 2 mM TCEP at 4℃ for 3h. Titrations were set up with MBP-eVP35_NTD_ protein in the syringe and MIB2_59-894_ protein in the cell. Different concentrations in the cell and injection were tested, with a reference performed using a power rate of 4 μcal/s. The resulting ITC data were processed and fit to a one-site-binding model to determine K_D_ (dissociation constant) using the ORIGIN 7.0 software.

### Fluorescence Polarization Assay (FPA)

FPA experiments were performed on a Cytation5 (Agilent Technologies) plate reader operating on Gen5 software. Fluorescein isothiocyanate (FITC)-eVP35_190-215_ or unlabeled peptides were purchased from Genscript, and MIB2_59-894_ was expressed and purified from *E. coli* as described previously. Excitation and emission wavelengths were set to 485 and 528 nm, respectively, with a bandpass of 20 nm. Sample read height was 7 nm and G factor was set to 1.26 using the autogain function within the Gen5 software. For binding experiments, a range of 0 – 15 µM MIB2_59-894_ was titrated into 2.5 nM FITC-eVP35_190-215_ or in buffer containing 20 mM HEPES (pH 7.0), 200 mM NaCl, and 2 mM TCEP. After 10 min of incubation, fluorescence polarization (FP) signals were read. The FP values were plotted against MIB2_59-894_ concentrations to fit the dissociation constant, *K*_D_, using GraphPad Prism software. For competition experiments, 2.5 nM FITC-eVP35_190-215_ was incubated with 2 µM MIB2_59-894_ the same buffer, and unlabeled eVP35 of mVP35 peptides at concentrations ranging from 0 - 500 µM with 5-fold dilution were loaded into a 96-well plate. After 10 min of incubation, FP signals were read. The FP values were plotted against eVP35 or mVP35 peptide concentrations to fit the dissociation constant, *Ki*, using GraphPad Prism software (La Jolla, CA).

### IFN-β reporter assay

IFN-β promoter reporter assays were performed in HEK 293T cells using a Dual-Luciferase Reporter assay (Promega). IFN-β was stimulated with RIG-I CARD domain in pCAGGS (50 ng) or low molecular weight Poly(I:C) (100 ng, invivogen), and they were co-transfected with WT or mutant eVP35 and/or MIB2 (all in pCAGGS) at concentrations indicated in the results, along with 200 ng of IFN-β promoter–firefly luciferase reporter plasmid and 20 ng of a constitutively expressed Renilla luciferase reporter plasmid. Transfections were done in suspension, using Lipofectamine 2000 and OptiMEM per the manufacturer’s recommendation. 48 h post-transfection, cells were harvested, and the Dual-Luciferase reporter assay (Promega) was used to measure luciferase levels. Firefly luciferase was normalized Renilla luciferase activity, the results were expressed as percent induction relative to the positive control. Graphpad Prism was used to analyze and present results. Error bars represent the s.d. of at least three independent experiments, in technical duplicate or triplicates each.

### NanoBRET Ubiquitination Assays

The system was established using the NanoBRET Ubiquitination Starter Kit (Promega ND2690). NanoLuc-CYLD was cloned into the pNLF1-N vector using XhoI and NotI restriction sites, and HaloTag-Ub was included in the kit. Subsequent experiments were ran using the NanoBRET Nano-Glo Detection System (Promega N1661). For ubiquitination experiments with transient expression, 1 × 10⁶ 293T cells were transfected in 6-well plates using TransIT LT1 transfection reagent (Mirus) with 2 μg of HaloTag-Ubiquitin and 0.02 μg NanoLuc-CYLD, with or without 0.1 μg eVP35 WT, eVP35 5A, or MIB2, followed by overnight incubation. On day 2, cells were washed with PBS, trypsinized, counted, and diluted in Opti-MEM (Gibco) with 4% FBS for a concentration of 2.2 × 10⁵ cells/mL. HaloTag NanoBRET 618 Ligand (Promega) was added to 400 μL of cells, and 90 μL per well (final concentration of 2.2 × 10⁴ cells/well) was plated in triplicate into white 96-well tissue culture plates (Corning), including background control wells without the ligand. Cells were incubated overnight at 37 °C with 5% CO₂. On day 3, NanoBRET NanoGlo (Promega) was prepared at 5× concentration in Opti-MEM (Gibco) and added before measuring NanoBRET signals using a Cytation5 microplate reader (Biotek) with a NanoBRET filter cube. Dual-filtered luminescence was collected using a 450/50 nm bandpass filter for donor emission (NanoLuc-CYLD) and a 610 nm long-pass filter for acceptor emission (HaloTag NanoBRET ligand). Readings were normalized to the HaloTag-Ub + NanoLuc-CYLD sample (set to 100%), and NanoBRET ratios were calculated as (acceptor channel/donor channel) × 1000. Data were analyzed and graphed using GraphPad Prism. Error bars represent the s.d. of at least three independent experiments, in technical triplicates each.

### TNFα stimulation

293T cells were transfected with either 0.5 μg or 2 μg of eVP35 WT or eVP35 5A using TransIT-LT1 transfection reagent (Mirus) according to the manufacturer’s instructions. After 24 hours, cells were treated with 20 ng/mL human TNFα (Thermofisher, 300-01A-100UG) and 0.5 μM BV6 (Invivogen, inh-bv6) for 2 hours. Cells were then harvested in Tritonx-100 buffer (25 mM Tris (pH 8), 150 mM NaCl, 1 mM EDTA, 1% TritonX-100, 10% glycerol) supplemented with protease and phosphatase inhibitors (cell signaling 5872), and an equal volume of 2x Laemmli loading dye was added. Whole-cell lysates were resolved by SDS-PAGE and analyzed by immunoblotting with the antibodies, as indicated in figures and figure legends.

### Minigenome Assays

We used the minigenome system as previously described.^37,38^ HEK293T cells were transfected with plasmids expressing viral proteins pCAGGS-eVP35 (125 ng), -eL (1000 ng), -eNP (125 ng), -eVP30(75 ng) along with pCAGGS-T7 polymerase (250 ng) and minigenome reporter plasmid (pM1-MG, 250 ng) that encodes for Renilla luciferase flanked by the minimally required cis-acting transcription and replication signals from the EBOV genome. A plasmid that expresses Firefly luciferase (2 5ng) was co-transfected and served as an internal control. MIB2 plasmids were co-transfected as indicated to determine how MIB2 impacts minigenome activity. Reporter activity was read 72 h post-transfection using the Dual-luciferase Reporter assay system (Promega). Renilla luciferase activity was normalized to Firefly luciferase values and plotted as minigenome activity.

### Cytopathic effect (CPE) assay

2 x 10^5^ Vero E6 cells were seeded in 12-well plates and incubated overnight. The media was removed, and cells were infected with rEBOV WT and rEBOV 5A (MOI 0.01) in 0.5 mL of inoculum for 30 minutes. The inoculum was removed, cells were washed once with PBS and 1 mL of DMEM 2% FBS was added. Images of wells were taken 7 dpi when CPE was evident using the brightfield channel on a EVOS FL microscope.

### Growth Kinetics Assay in Vero E6 Cells

Vero E6 cells (2 × 10⁵ cells/well) were seeded into seven 24-well plates 12 hours prior to infection. Cells were infected in triplicate with either wild-type (WT) EBOV or the VP35 NNLNS mutant EBOV at a multiplicity of infection (MOI) of 0.01 using 250 µL of viral inoculum per well. After a 30-minute adsorption period at 37°C, the inoculum was removed and cells were washed once with phosphate-buffered saline (PBS). Fresh 500 µL of Dulbecco’s Modified Eagle Medium (DMEM) supplemented with 2% fetal bovine serum (FBS) was added to each well. At indicated time points (daily for 6 days), 500 µL of supernatant was harvested from each well, and viral titers were determined by focus-forming assay on Vero E6 cells.

### Mass photometry

Clean coverslips were assembled into flow chambers and mounted on the slide with an oil drop per the manufacturer’s instructions. Immediately prior to mass photometry measurements, protein stocks were diluted directly in stock buffer (20 mM HEPES, 200 mM NaCl, 2 mM TECEP, at pH 8.0). To find focus, 15 – 18 µL fresh buffer was first flowed into the chamber, the focal position was identified and secured in place with an autofocus system. For each acquisition, 2 – 5 µL of protein was added to the buffer in the chamber (for a final working concentration of 10 - 20 nM), mixed, and measurements acquired for 60 seconds. Each sample was measured at least three times independently. The calibration protocol included measurement of two proteins thyroglobulin (660 kDa), and b-amylase (monomer of 55 kDa and dimer of 110 kDa). Each MP calibration experiment was analyzed using DiscoverMP. The mean peak contrast was determined in the software using Gaussian fitting.

### Size-exclusion chromatography coupled to multi-angle light scattering (SEC-MALS)

SEC-MALS experiments were carried out using buffer containing 20 mM HEPES, 150 mM NaCl, and 2 mM TCEP, at pH 8.0. Protein concentration between 1.2 – 3 mg/mL were used. Purified MBP-VP35_1-215_, MIB2_59-894_, or a complex of the two were autoinjected on a DAWN-HELEOS II detector (Wyatt Technologies) coupled to a Superdex 75 Increase 5/150 GL column (GE Healthcare). The acquired data from UV, MALS, and RI detectors were analyzed to determine weight-averaged molecular mass (Mw) of the proteins using ASTRA 6 software (Wyatt Technologies). Protein concentrations were determined using the refractive index measured by an Optilab T-rEX (Wyatt Technologies) and dn/dc = 0.185 ml^3^g^-1^. Bovine serum albumin (ThermoFisher) was used as a calibration standard.

### Quantification and Statistical Analyses

ITC data were fitted to a one-site-binding model to determine KD (dissociation constant) using the ORIGIN 7.0 software. All other statistical analysis was performed using GraphPad Prism 7. All statistical details and the appropriate definition of parameters can be found in the figure legends and text. Data points were considered significantly different if the p-value was < 0.05.

## Acknowledgements

EBOV pCAGGS support plasmids and p15AK EBOV Mayinga were kind gifts from Drs. Elke Muhlberger and Heinz Feldmann, respectively. We thank R. Andrews and H. Kohlmiller for their assistance with hybridoma generation and screening. We thank L. Wheat for coordinating studies between WUSM and external groups. This work was supported by NIH grants P01120943 (R.A.D., C.F.B., D.W.L., and G.K.A.), U19AI109664 (C.F.B.), U19AI109945 (C.F.B.), P50GM082250 (N.J.K.), U19AI106754 (N.J.K.), R01AI120694 (N.J.K.), P01AI063302 (N.J.K.).

## Author contributions

GU, GKA, and DWL conceived the overall project. All authors were integral to the design and execution of the study. GU, GKA, and DWL wrote the initial draft with significant input from all authors.

## Conflicts of interest

N.J.K. has received research support from Vir Biotechnology, F. Hoffmann-La Roche, and Rezo Therapeutics. N.J.K. has a financially compensated consulting agreement with Maze Therapeutics. N.J.K. is the President and is on the Board of Directors of Rezo Therapeutics, and he is a shareholder in Tenaya Therapeutics, Maze Therapeutics, Rezo Therapeutics, GEn1E Lifesciences, and Interline Therapeutics.

