## Supplemental file for "Ebola virus VP35 NNLNS motif modulates viral RNA synthesis and MIB2-mediated signaling"

**Inventory of Supplemental Information**

- **Supplemental Data**
  - **Figure S1, Related to Figure 2.** eVP35 binds MIB2
  - **Figure S2, Related to Figure 3.** eVP35 linker residues 201-205 are critical for MIB2 binding
  - **Figure S3, Related to Figure 4.** eVP35 inhibits MIB2-mediated IFN induction
  - **Figure S4, Related to Figure 5**. eVP35 $^{201}$NNLNS$^{205}$ motif contributes to EBOV RNA synthesis
  - **Figure S5, Related to Figure 5**. VP35-MIB2 interaction with VP35 from different filoviruses

FIGURE CAPTIONS

**Figure S1. eVP35 binds MIB2**

(**A**) Representative size-exclusion chromatogram of MBP-tagged eVP35$_{1-215}$ (black), MIB2$_{59-894}$ (blue), and complex (red). The complex was formed by incubating both proteins at room temperature for 10 minutes. Shown also are Coomassie-stained gels of fractions collected from each SEC run. (**B**) Domain architecture of MIB2 constructs tested to bind eVP35. Purified MBP-eVP35 or MBP alone were immobilized on amylose resin prior to the addition of $MIB2_{1-1013}$ (**C**) $MIB2_{\Delta RING}$ (**D**) $MIB2_{59-894}$ (**E**). Shown are the Coomassie-stained SDS-PAGE gels of each sample. Each gel is representative of at least 2 independent replicates. (Input, marker (M), flow-through (FT), washes (W1, W2, W3, W4, and W5), beads after washes (B) are indicated above each lane.

**Figure S2. eVP35 linker residues 201-205 are critical for MIB2 binding**

(**A**) A list of VP35 constructs used to determine the minimum region of eVP35 required to interact with $MIB2_{1-894}$. Pull-down assays were used to test the binding between MBP-tagged VP35 constructs and $MIB2_{58-894}$. (**B-F**) show examples of Coomassie-stained SDS-PAGE gels used to analyze the pull-down assays. (**G**) Mass photometer profiles of MBP-$VP35_{1-215}$ WT (blue) and MBP-$VP35_{1-215}$ 5A (green). Indicated are calculated masses, standard error, and number of counts for each mass. Both proteins are shown to be tetrameric. (**H**) Representative ITC raw data and corresponding binding isotherms show a loss of eVP35 binding to MIB2 when residues $^{201}$NNLNS$^{205}$ are mutated. Each experiment was done with $MIB2_{59-894}$ in the cell. From left to right: MBP-$VP35_{1-215}$ WT in the injection. $K_D$ of 5.4 ± 2.5 μM, N = 0.9 ± 0.2, ∆H = -22.0 ± 0.2 kcal/mol; MBP-$VP35_{1-215}$ 5A in the injection. $K_D$ = N.D, N = N.D.; MBP-$VP35_{1-215}$ 3A in the injection. $K_D$ = N.D, N = N.D.; MBP-$VP35_{1-215}$ 2A in the injection. $K_D$ = N.D, N = N.D. The data shown are representative of at least 3 independent experiments. N.D, not detected. 3A = residues $^{203}$LNS$^{205}$ mutated to A, 2A = residues $^{201}$NN$^{202}$ mutated to A.

**Figure S3. eVP35 inhibits MIB2-mediated IFN induction**

(**A**) An IFN-β luciferase reporter assay when IFN production was induced by MIB2 WT or MIB2 RINGm. WT or RINGm (100 or 500 ng) was co-transfected with IFN-β firefly reporter plasmid, and luciferase activity was measured 48h post-transfection. (**B** and **C**) IFN-β luciferase assay when IFN production was co-stimulated by CARD domain (50 ng) and MIB2 WT or MIB2 RINGm (500 ng), in the presence of 500 ng of eVP35 WT (blue) or eVP35 5A mutant. (**D**) An IFN-β luciferase assay in the presence of empty vector (pCAGGS) or 1000 and 2000 ng of eVP35 WT (blue), 5A (green), KRA (orange), or 5A/KRA (cyan) mutant. IFN was induced with 50 ng CARD domain (black), and 48h post-transfection IFN-β activity was assessed by measuring luciferase activity. Statistical comparisons are between WT eVP35 and each mutant at the same concentration. The statistical significance is as indicated: not significant: n.s., *$p < 0.05$, **$p < 0.01$, ***$p < 0.001$, and ****$p < 0.0001$. (**E**) 293T cells were seeded for 24h and treated with 0.5 μM BV6 and indicated amounts of TNFα for 2h. WB was used to analyze CYLD protein levels.

**Figure S4. eVP35 linker residues 201-205 contribute to EBOV RNA synthesis**

**(A)** eVP35 WT (blue), eVP35 5A mutant (green), or a mixture of the two at indicated percentages was cotransfected with the rest of the plasmids required to reconstitute the EBOV RNA polymerase complex (L, NP, VP30) along with a plasmid encoding the Renilla luciferase minigenome RNA and a firefly luciferase expression plasmid, which served as a control for transfection efficiency. Relative activity was determined by setting WT VP35 Renilla luciferase to 100% activity and normalizing the rest of the conditions against it. (**B**) Infection efficiency was assessed in WT 293T cells using recombinant WT EBOV (rEBOV) and EBOV carrying the NNLNS substitution in VP35 (rEBOV-5A), both encoding ZsGreen as an infection marker. Cells were infected at an MOI of 0.1 to limit infection to a single event per cell. After inoculation, cells were washed twice with PBS and incubated. At 20 hours post-infection (hpi), cells were fixed with formalin and stained with Hoechst 33342 to visualize nuclei. ZsGreen-positive cells were quantified above a defined detection threshold. Infection efficiency was calculated as the percentage of infected cells per total cells and normalized to the mean infection level of WT virus. Eight replicates were performed per condition, with individual values, means, and standard deviations shown. Statistical analysis was conducted using one-way ANOVA with multiple comparisons to WT; significance is indicated (**** = $P < 0.0001$). (**C**) Growth kinetics of rEBOV-WT and rEBOV-5A in Vero E6 cells. Vero E6 cells were infected in triplicate at a multiplicity of infection (MOI) of 0.01 with either virus. Supernatants were collected daily for 6 days post-infection, and viral titers were determined by focus-forming assay on Vero E6 cells. Titers are expressed as $\log_{10}$ focus-forming units per milliliter (FFU/mL) and represent the mean ± standard deviation (SD) of three independent replicates. (**D**) Increasing amounts of MIB2 WT or MIB1 plasmid (125, 500 ng) were co-transfected with the plasmids required to reconstitute the EBOV RNA polymerase complex in the presence or absence of eVP35 WT. (**C**) Similar to B, MIB2 WT or MIB2 RINGm (125 or 500 ng) was added in the minigenome assay, either with eVP35 WT (blue) or eVP35 5A (green). Statistical comparisons are between MIB2 WT and MIB2 RINGm of the same concentration. Protein expression was validated by WB. The statistical significance is as indicated: not significant: n.s., *$p < 0.05$, **$p < 0.01$, ***$p < 0.001$, and ****$p < 0.0001$. (**D**) eVP35 WT (blue) and eVP35 5A (green) are ubiquitinated in the presence of MIB2 WT but not MIB2 RINGm or MIB2 ΔRING. Flag-eVP35 or Flag-eVP35 5A and HA-Ubiquitin were transfected in the presence or absence of MIB2 WT, MIB2 RINGm, or MIB2 ΔRING, and immunoprecipitation with HA magnetic beads was performed. WB was used to analyze the interaction. Ubiquitinated VP35 is indicated with arrows.

**Figure S5. VP35-MIB2 interaction with VP35 from different filoviruses.** (**A**) Structure of EBOV VP35-L complex, adopted from Yuan et. al. 2022 cyro-EM structure, PDB file 7YES. The data file was downloaded from the Protein Data Bank, and Pymol was used to adapt the colors and annotations. Light purple = polymerase L, grey and green = VP35 tetramer. eVP35 residues $^{201}$NNLNS$^{205}$, K319, and R322 are labeled. Also indicated are the shortest distances between eVP35 NNLNS residues and L residues F786, R400, and A398. Note that our studies use eVP35 from the Mayinga '76 strain, which has the sequence $^{201}$NNLNS$^{205}$, while this structure corresponds to eVP35 from the Makona strain which has the sequence $^{201}$NNLDS$^{205}$ instead. (**B**) MIB2 interacts with VP35 from different filoviruses. HA-tagged VP35 was co-expressed with Flag-tagged MIB2, and a co-IP experiment was performed with HA magnetic beads as described above. EBOV = Ebola virus, MARV = Marburg virus, Bat = *Myotis myotis* bat species. (**C**) MARV $VP35_{FL}$ or (**D**) EBOV $VP35_{1\text{-}215}$ were immobilized on amylose resin before the addition of purified $MIB2_{59\text{-}894}$. After 5 washes, the complex was eluted with 1% maltose. Shown are the Coomassie-stained SDS-PAGE gels of each sample. Lane labels: FT = flow-through, W = wash, B = beads, E = elution. (**E**) Competition of unlabeled EBOV $VP35_{190\text{-}205}$ (black) or MARV $VP35_{179\text{-}204}$ (blue) peptides with 2.5 nM FITC-$eVP35_{190\text{-}215}$ in the presence of 2 μM $MIB2_{59\text{-}894}$ resulted in the indicated competition curves. $K_{D,App}$: 4.71 ± 2.20 μM for

EBOV $VP35_{190-205}$, N.D for MARV $VP35_{179-204}$. (**F**) Sequence alignment of the linker region containing the NNLNS motif (red box) of EBOV (Mayinga ’76, Genbank accession CAA43578.1, Makona ’14 ALX31317.1), MARV (Musoke ’08 CAA78115.1), and a VP35 homologue encoded in the bat Myotis myotis genome (AXC07973.1). Amino acid sequences for each VP35 were aligned using Clustal omega and imported into Jalview, and the residues are colored by percentage identity conservation. The alignment is numbered with respect to the EBOV Mayinga ’76 sequence that was used in this study.

**Supplemental figure 1**

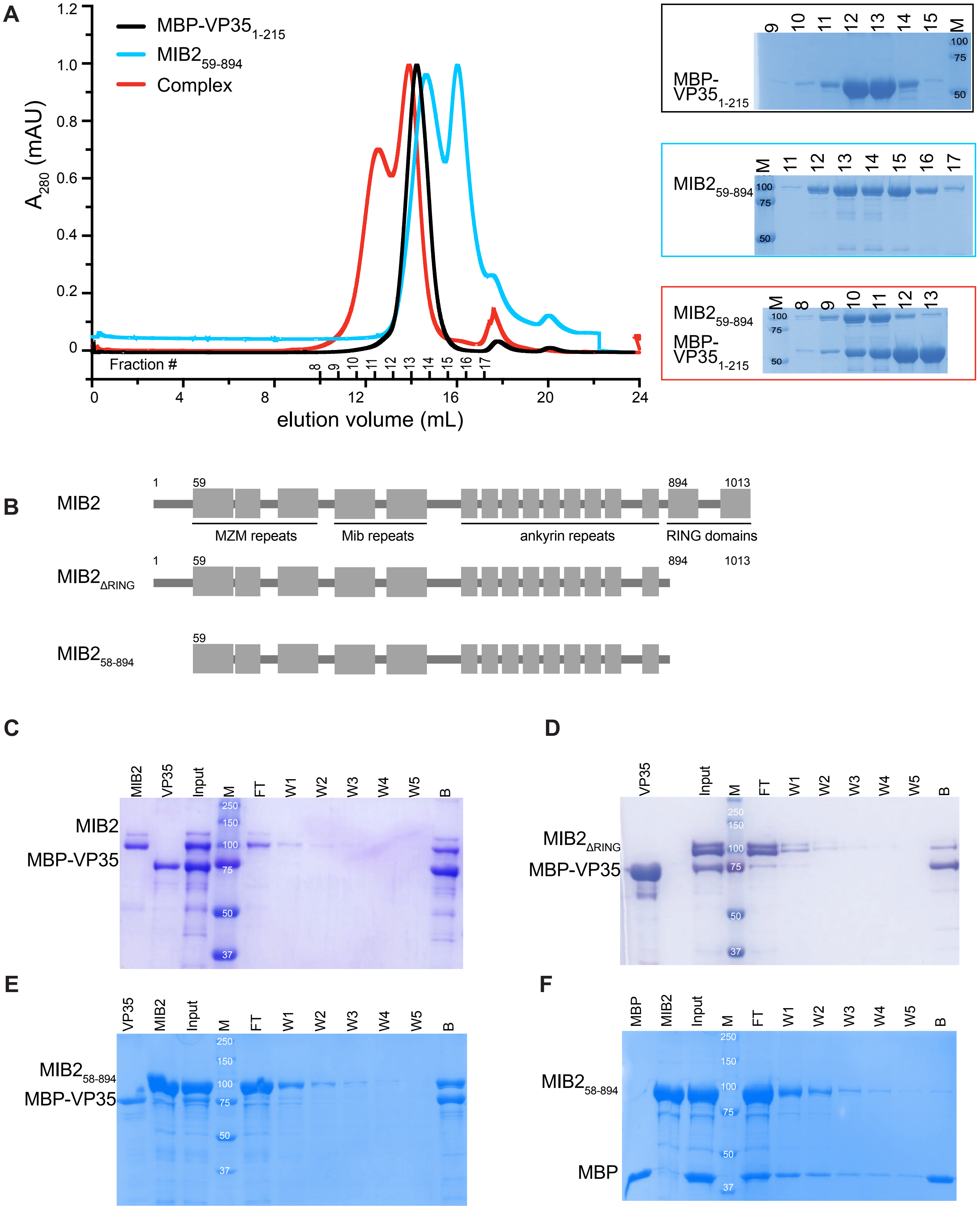

### Supplemental figure 2

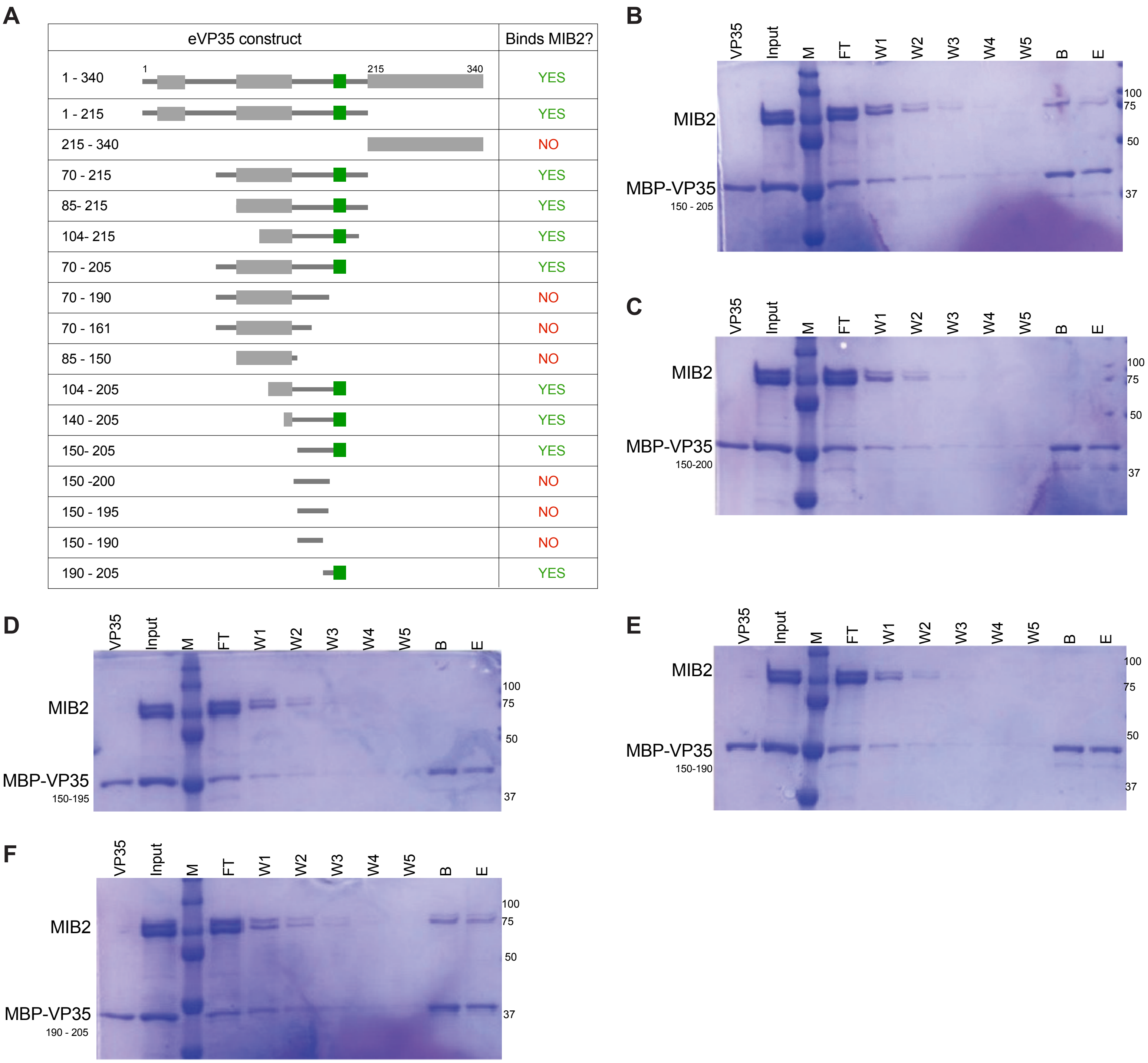

#### Supplemental figure 2 continued

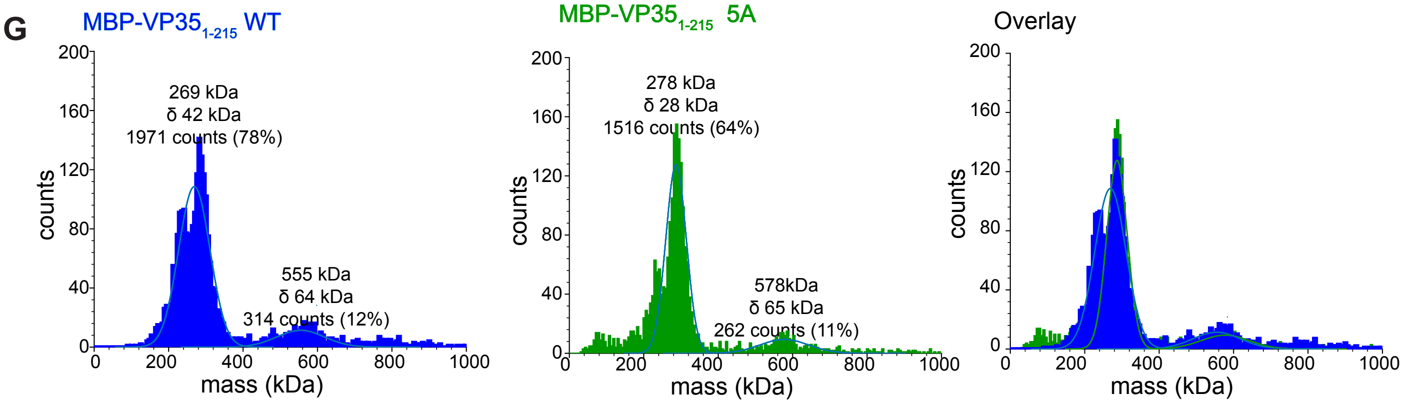


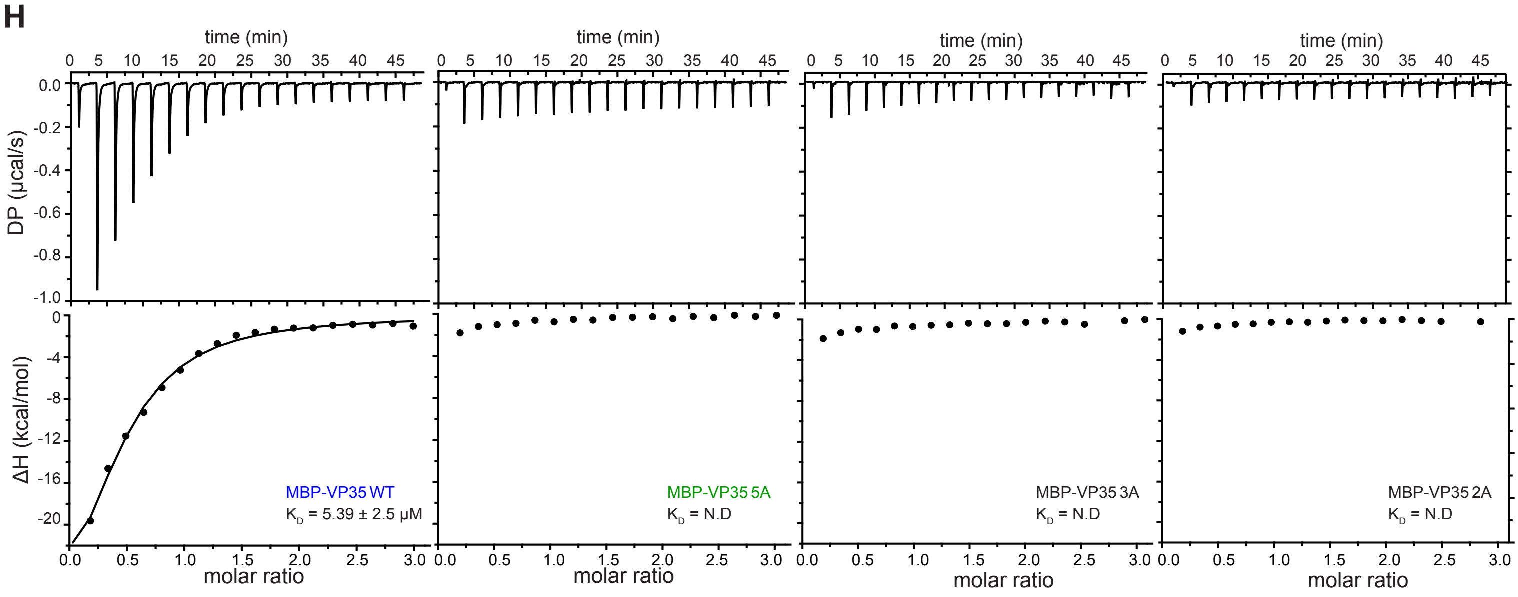

### Supplemental figure 3

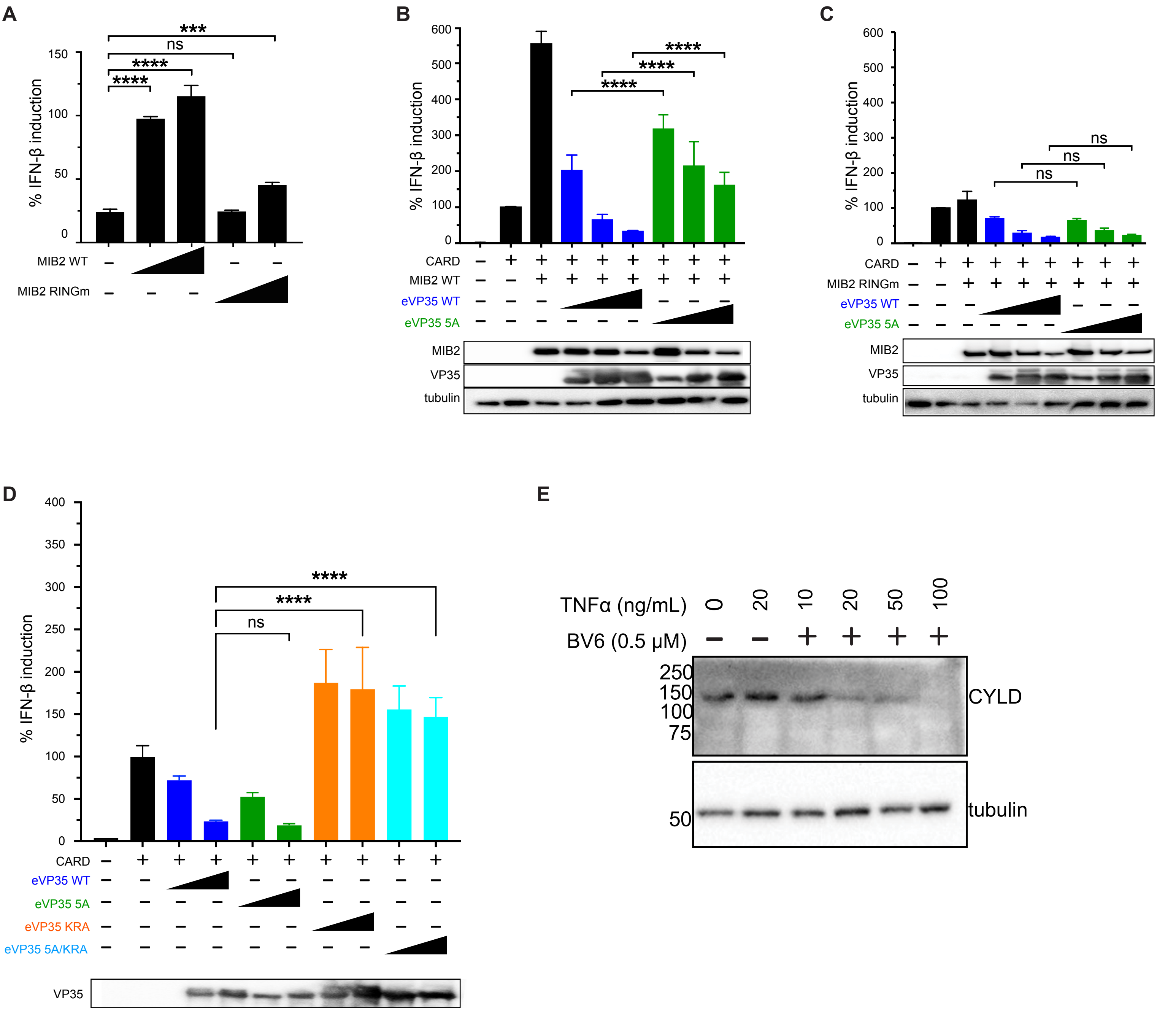

#### Supplemental figure 4

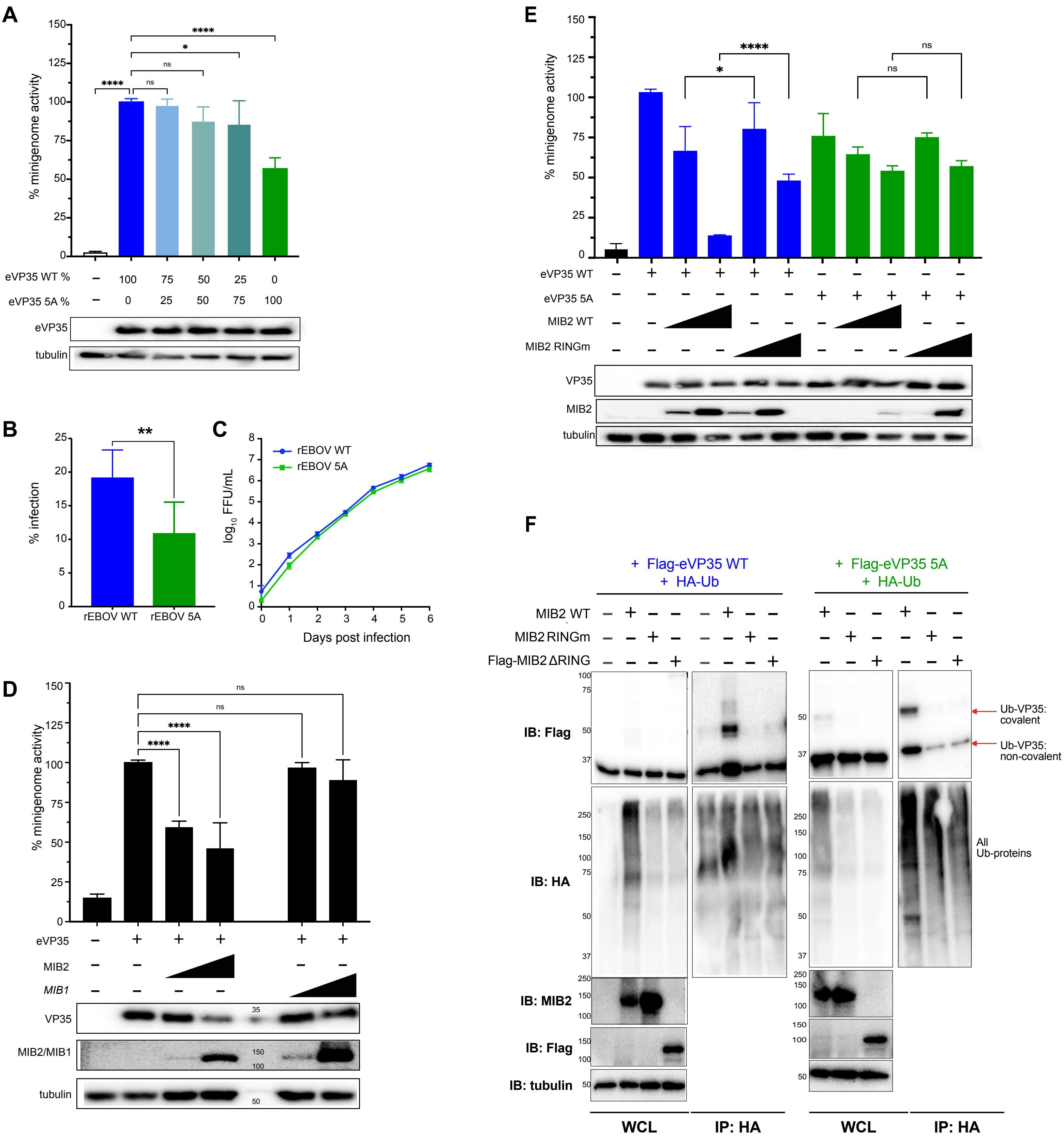

### Supplemental figure 5

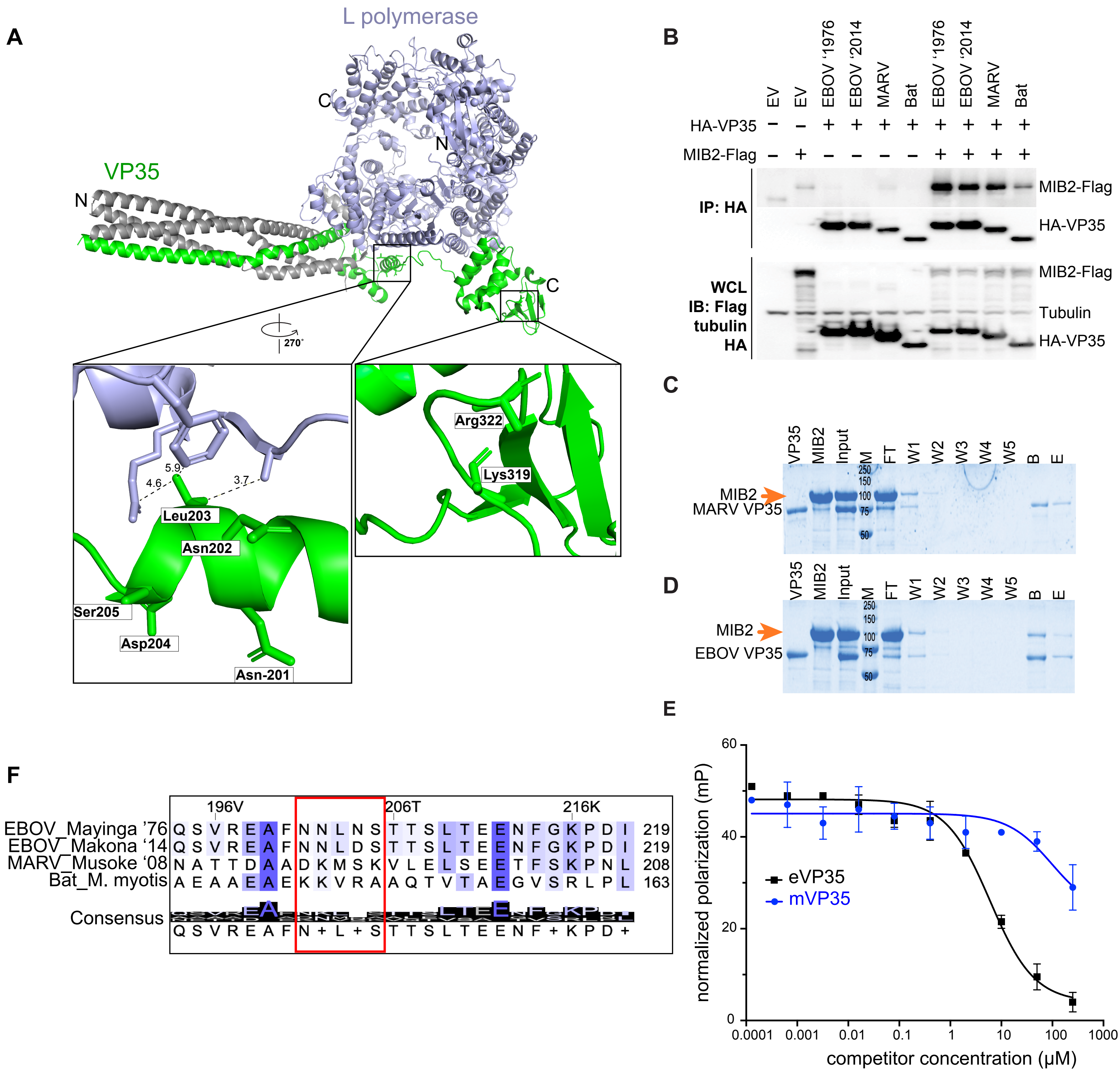
